# ACSM1 and ACSM3 regulate prostate cancer fatty acid metabolism to promote tumour growth and constrain ferroptosis

**DOI:** 10.1101/2022.10.13.511039

**Authors:** Raj Shrestha, Zeyad D. Nassar, Adrienne R. Hanson, Richard Iggo, Scott L. Townley, Jonas Dehairs, Chui Yan Mah, Madison Helm, Mohammadreza Ghodsi, Marie Pickering, Matthew J. Watt, Lake-Ee Quek, Andrew J. Hoy, Wayne D. Tilley, Johannes V. Swinnen, Lisa M. Butler, Luke A. Selth

## Abstract

Prostate tumours are highly reliant on lipids for energy, growth and survival. Activity of the androgen receptor (AR) is associated with reprogramming of lipid metabolic processes in prostate cancer, although the molecular underpinnings of this relationship remain to be fully elucidated. Here, we identified Acyl-CoA Synthetase Medium Chain Family Members 1 and 3 (ACSM1 and ACSM3) as AR-regulated mediators of prostate cancer metabolism and growth. ACSM1 and ACSM3 are upregulated in prostate tumours compared to non-malignant tissues and other cancer types. Both enzymes enhanced proliferation and protected PCa cells from death *in vitro*, while silencing ACSM3 led to reduced tumour growth in an orthotopic xenograft model. We show that ACSM1 and ACSM3 are major regulators of the PCa lipidome and enhance energy production via fatty acid oxidation. Metabolic dysregulation caused by loss of ACSM1/3 led to mitochondrial oxidative stress, lipid peroxidation and cell death by ferroptosis. Conversely, over-expression of ACSM1/3 enabled PCa cells to survive toxic doses of medium chain fatty acids and promoted resistance to ferroptosis-inducing drugs and AR antagonists. Collectively, these studies uncover a new link between AR and lipid metabolism and position ACSM1 and ACSM3 as key players in prostate cancer progression and therapy resistance.

## INTRODUCTION

Cancer cells have different metabolic requirements compared to normal cells, as they are highly proliferative and need to survive in a transformed microenvironment characterised by limited nutrient supply. Altered lipid metabolism represents one of the most common metabolic adaptations exhibited by cancer cells^1^. For example, cancer cells exhibit increased *de novo* synthesis and uptake of fatty acids (FAs), which are used for membrane biogenesis, energy production via β-oxidation, storage as triglycerides, and protein modification^2^. Additionally, cancer cells actively stimulate mobilisation and release of stored lipids from adipocytes in the tumour microenvironment^2^.

Prostate cancer (PCa) is characterised by and is highly dependent on reprogramming of lipid metabolic pathways^3^, a phenomenon that is heavily influenced by the androgen receptor (AR) signalling axis^4^. AR, a ligand (androgen)-activated transcription factor, is the primary oncogenic driver of PCa and the major therapeutic target in advanced and metastatic disease^5^. Failure of AR-targeted therapies, such as androgen deprivation therapy (ADT) and AR antagonists (e.g. enzalutamide), results in an aggressive and lethal form of the disease termed castration-resistant prostate cancer (CRPC). AR indirectly influences key regulators of lipid metabolism by enhancing the expression and activity of sterol regulatory element binding proteins (SREBPs)^4^, transcription factors with a fundamental role in activating a lipogenic transcriptional program. Additionally, AR directly regulates the expression of genes encoding proteins involved in lipid synthesis, uptake and storage^4, 6^, one prominent example being fatty acid synthase (FASN)^7^. Given the essential role of FAs/lipids in enhancing the growth of PCa and the emerging realisation of their widespread and intricate interplay with AR signalling, there is increasing interest in therapeutic targeting of lipid metabolic pathways^8–10^.

The first step in utilisation of FAs - either those synthesised *de novo,* hydrolysed from intracellular glycerolipids, or taken up from exogenous sources - is ATP-dependent generation of a thioester bond with coenzyme A (CoA) to form fatty acyl-CoA esters. This thioesterification reaction is catalysed by a large family of enzymes called acyl-coenzyme A (acyl-CoA) synthetases (ACSs; also known as acyl-CoA ligases) and yields substrates for β-oxidation, protein acylation and lipid synthesis^11^. The carbon chain length of FAs varies from 2 to >30, which dictates processing by distinct sub-families of ACSs with selectivity for short-chain (C2–C4, ACCSs), medium-chain (C4–C12, ACSMs), long-chain (C12–C22, ACSLs), bubblegum (C14-C24, ACSBG), and very long-chain (C18–C26, annotated as solute carrier family 27A) fatty acids^12^. Despite their central roles in cellular lipid metabolism, the normal and pathophysiological functions of many ACS enzymes, particularly the ACSMs, are poorly understood.

Here, we identify ACSM1 and ACSM3 as previously unrecognised AR-regulated enzymes that are highly expressed in the malignant prostate. Functional assays coupled with lipidomics and metabolomics revealed that ACSM1/3 play critical roles in PCa metabolism, growth and survival, with implications for tumour response to AR antagonists and ferroptosis-inducing drugs. Collectively, these data identify a new mechanism by which AR influences PCa lipid metabolism and reveal medium chain fatty acid activation by ACSM1/3 as an unexplored therapeutic vulnerability.

## RESULTS

### ACSM1 and ACSM3 are AR target genes in prostate cancer

To identify novel downstream effectors of AR in PCa, we first compiled a list of 178 genes encoding factors with a putative role in lipid metabolism (Table S1). Subsequently, published data^13–17^ was interrogated to filter the gene set by 3 criteria (Figure 1A): 1) evidence of regulation by androgen treatment in models of PCa; 2) evidence of proximal AR DNA binding proximal to transcriptional start sites (TSSs); and 3) evidence of dysregulated expression in clinical PCa. This data mining effort revealed 17 genes of interest: *ACACA*, *ACOT2*, *ACSL3*, *ACSM1*, *ACSM3*, *ELOVL5, ELOVL7*, *FASN, HACL1, HSD17B4, MAPKAPK2, MMAA, MORC2, PCCB, PCTP, PRKAB2* and *PTGR2* (Figure 1A; Table S1). The identification of known androgen-regulated genes (e.g. *ACACA*^6^, *ACSL3*^18^, *ELOVL5*^19^ and *FASN*^20^) validated our approach. We focussed our attention on *ACSM1* and *ACSM3* (<u>A</u>cyl-<u>C</u>oA <u>S</u>ynthetase <u>M</u>edium Chain Family Members 1 and 3), genes that encode enzymes reported to catalyse the activation of medium chain fatty acids (MCFAs), since their potential roles in PCa are entirely unknown. AR-mediated upregulation of *ACSM1* and *ACSM3* was confirmed by treatment of LNCaP and VCaP cells with the potent androgen dihydrotestosterone (DHT), an effect that was reversed by co-treatment with the AR antagonist enzalutamide (Enz; Figure 1B-C). These findings were reinforced by transfecting LNCaP or VCaP cells with an AR siRNA, which reduced both expression and protein levels of *ACSM1* and *ACSM3* (Figures 1D-E). In support of the *in vitro* data, *ACSM1* and *ACSM3* expression decreased following ADT in patient tumours (Figure 1F).

**Figure 1.**
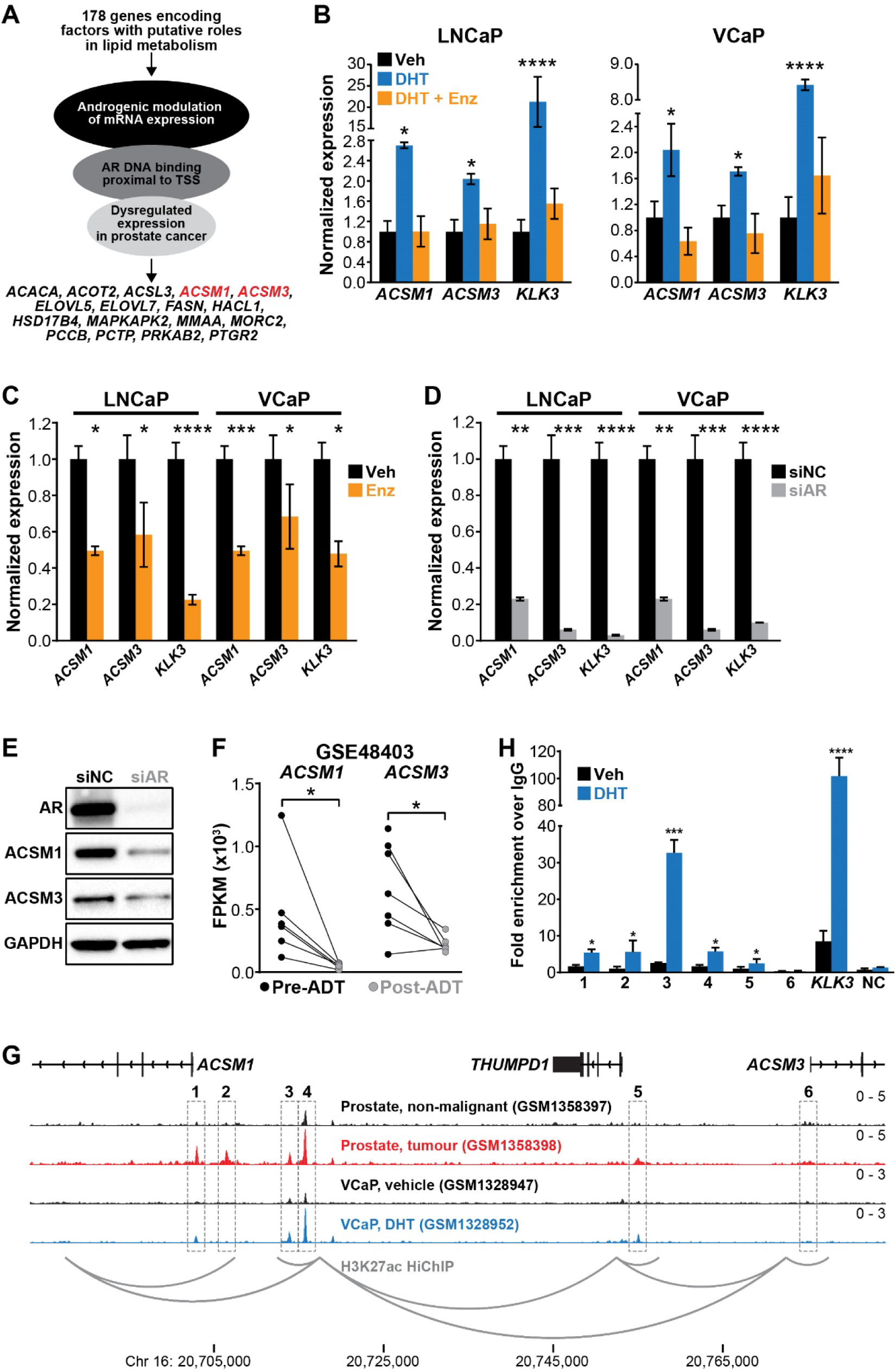
*ACSM1* and *ACSM3* are AR target genes in prostate cancer. **(A)** Schematic illustration of strategy to identify novel AR-regulated lipid metabolic genes. **(B-D)** Expression of *ACSM1* and *ACSM3* in response to: **(B)** 24 h of treatment with DHT (10 nM) or DHT + enzalutamide (10 µM) (DHT+Enz) in cells grown in charcoal-stripped serum; **(C)** 48 h of treatment with enzalutamide (Enz) in cells grown in full serum; and **(D)** 72 h of transfection with AR-specific siRNA (siAR). Gene expression was normalised to *GAPDH;* vehicle (Veh) or negative control siRNA (siNC) were set to 1; error bars are SEM. Differential expression was evaluated using unpaired t tests. **(E)** ACSM1 and ACSM3 protein levels in response to siAR treatments were measured by immunoblotting in LNCaP cells. GAPDH was used as a loading control. **(F)** *ACSM1* and *ACSM3* mRNA expression in prostate tumours pre- and post- androgen deprivation therapy (ADT)^21^. A Wilcoxon matched-pairs signed rank test was used to compare expression in the groups. FPKM, fragments per kilobase of exon per million mapped reads. **(G)** ChIP-seq data showing AR DNA binding near the *ACSM1* and *ACSM3* gene in non-malignant and prostate tumour samples^13^ and VCaP cells^22^ and H3K27ac HiChIP data from LNCaP cells^23^. The grey dotted box indicates AR binding peaks. **(H)** ChIP-qPCR analysis of 6 putative AR binding sites (1-6; shown below the ChIP-seq tracks in G). Data is represented as fold-change over an IgG control. Error bars represent SEM. For all statistical tests: *, p < 0.05; **, p < 0.01; ***, p < 0.001; ****, p < 0.0001.

As part of our gene filtering criteria, we had noted putative AR binding sites proximal to the *ACSM1* and *ACSM3* TSSs in ChIP-seq data from prostate tumours and patient-matched normal specimens, some of which were linked by histone H3 lysine 27 acetylation (H3K27ac)-associated chromatin looping events (Figure 1G). The AR binding events were also induced by androgen treatment in AR ChIP-seq data from the VCaP cell line model, supporting their functional relevance (Figure 1G). We validated androgen-regulated AR binding at 5 out of 6 of these loci by ChIP-qPCR in LNCaP cells (Figure 1H). Collectively, these data suggest that AR directly regulates the expression of *ACSM1* and *ACSM3*.

### ACSM1 and ACSM3 are highly expressed in the malignant prostate

To assess the clinical relevance of ACSM1 and ACSM3 in PCa, we examined a series of published clinical datasets. Both genes were upregulated in PCa compared to non-malignant prostate tissues in three large cohorts, two of which (TCGA and CPGEA) represent patient-matched normal:tumour pairs (Figure 2A). Moreover, in multiple proteomic datasets, ACSM3 levels were elevated in cancer compared to non-malignant tissues (Supplementary Figure S1; ACSM1 was not evaluated in these studies). To extend upon these *in silico* analyses, we used immunohistochemistry to measure ACSM1 and ACSM3 protein expression in a tissue microarray containing non-malignant and tumour tissues from 160 men. Both proteins were detectable in all samples: ACSM1 is ubiquitously expressed in epithelia and stroma, whereas ACSM3 expression is restricted to epithelial cells (Figure 2B, right). Consistent with the public datasets, levels of both proteins were significantly higher in tumour sections and associated with increasing Gleason grade (Figure 2B, left; representative images shown on right).

**Figure 2.**
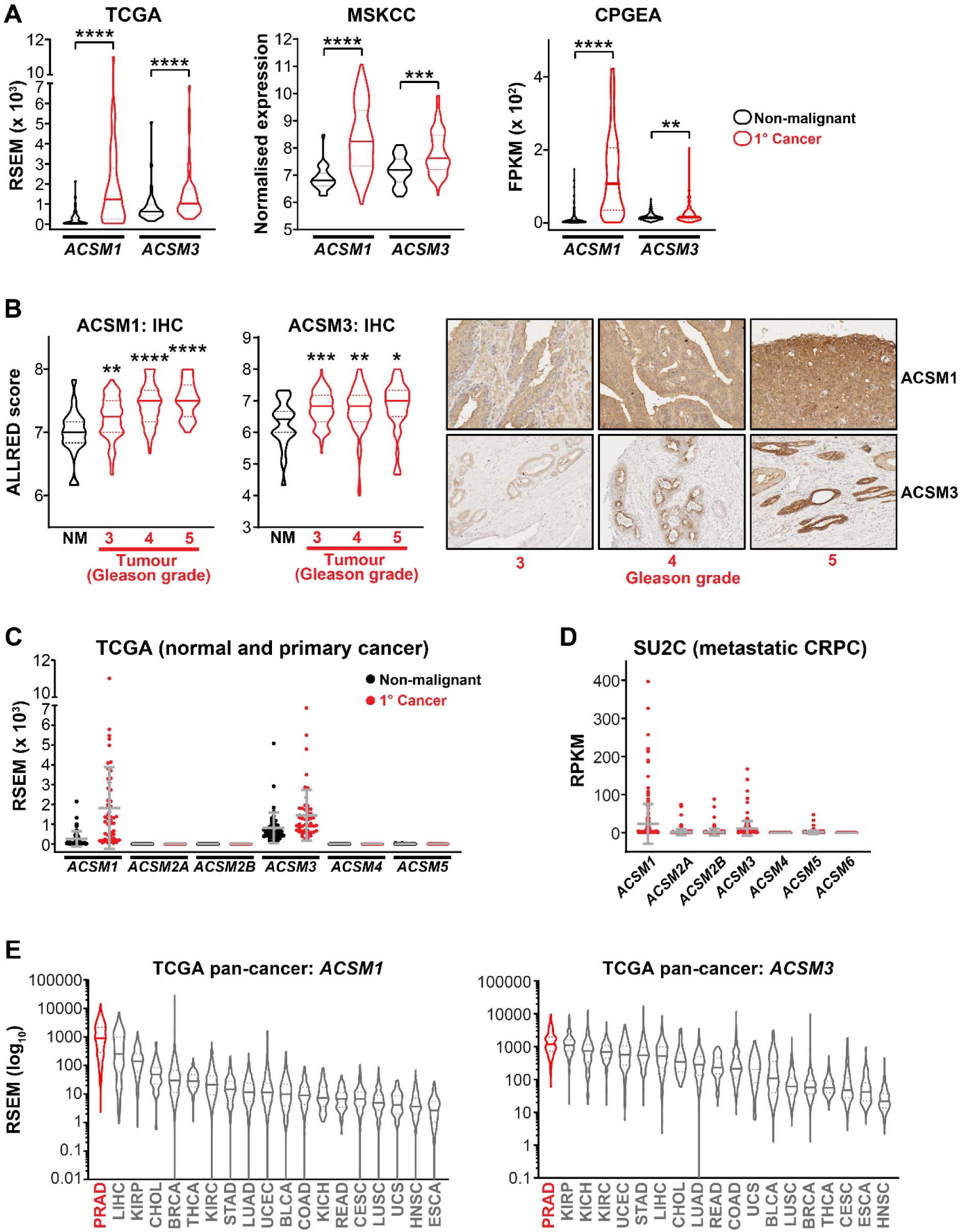
ACSM1 and ACSM3 are over-expressed in prostate cancer. **(A)** *ACSM1* and *ACSM3* expression is elevated in primary prostate cancer (1° Cancer). The TCGA dataset comprises 52 patient-matched non-malignant and cancer samples. Violin plots show minimum and maximum (bottom and top lines, respectively) and mean (line within the boxes) values. Paired (TCGA and CPGEA) or unpaired (MSKCC) t tests were used to compare expression in non-malignant versus cancer tissues. RSEM, a quantitative value produced by RNA-Seq by Expectation Maximization; FPKM, fragments per kilobase of exon per million mapped reads. **(B)** ACSM1 and ACSM3 protein levels are associated with prostate cancer grade, as determined by IHC. Violin plots on left show minimum and maximum (bottom and top lines, respectively) and mean (line within the plot) values. One-way ANOVA was used to compare primary Gleason grade groups with non-malignant tissues. Representative IHC images are shown on the right (scale bars represent 50 μm). **(C)** Expression of ACSM family members in normal prostate and primary tumours in the TCGA cohort. **(D)** Expression of ACSM family members in metastatic CRPC in the SU2C cohort. RPKM, reads per kilobase of exon per million mapped reads. **(E)** Pan-cancer expression of *ACSM1* and *ACSM3* using a unified TCGA dataset ^24^. For all statistical tests: *, p < 0.05; **, p < 0.01; ***, p < 0.001; ****, p < 0.0001.

The ACS medium chain family is comprised of 7 genes: *ACSM1*, *ACSM2A*, *ACSM2B*, *ACSM3*, *ACSM4*, *ACSM5* and *ACSM6*. To assess if other family members have a function in PCa, we interrogated RNA-seq data from the TCGA (non-malignant and primary cancer) and SU2C (metastatic CRPC) cohorts. *ACSM1* and *ACSM3* are the only family members expressed at an appreciable level in the normal prostate and primary tumours (Figure 2C). Although *ACSM2A*, *ACSM2B* and *ACSM5* are detectable in a subset of metastatic CRPC tumours, *ACSM1* and *ACSM3* are expressed at significantly higher levels (p < 0.0001 compared to all other genes; Figure 2D). We then evaluated the pan-cancer distribution of *ACSM1* and *ACSM3* expression using a unified version of the TCGA dataset in which data from distinct tumour types can be compared after correction for study-specific biases^24^. Notably, PCa exhibits the highest expression of both genes compared to 18 other cancer types (Figure 2E).

### ACSM1 and ACSM3 promote the growth of prostate cancer

Elevated expression of ACSM1 and ACSM3 in prostate tumours compared to benign tissues and other cancer types likely indicate important functions of these proteins in PCa growth and progression. To evaluate this hypothesis, we undertook functional studies in models of PCa. First, we examined expression of both proteins in a panel of cell lines: ACSM1 was robustly expressed in all 11 models tested, whereas the expression of ACSM3 was more variable and completely absent in the AR-negative cell lines (PC3 and DU145) (Figure 3A). Transient knockdown using two distinct siRNAs for each factor, both of which were highly effective at reducing cellular protein levels (Figure 3B), was used to determine whether ACSM1 and/or ACSM3 contribute to PCa growth. Silencing either protein markedly suppressed the proliferation of cell line models of castration-sensitive (LNCaP, VCaP) and castration-resistant (22Rv1, 49F^ENZR^) disease (Figure 3C). ACSM1 knockdown also caused inhibition of PC3 growth but as expected ACSM3-specific siRNAs had no effect in this ACSM3-negative line (Figure 3C). Concomitant knockdown of both ACSM1 and ACSM3 did not elicit a stronger growth inhibitory phenotype than either factor alone (Figure 3C), suggesting redundancy in their function. The relevance of ACSM1 and ACSM3 in PCa tumourigenesis and growth was further demonstrated using colony formation and 3D spheroid assays, the latter utilised because it more closely mimics *in vivo* conditions than 2-dimensional cell culture. In both assays, loss of ACSM1 and ACSM3 had profound tumour suppressive effects (Figure 3D-E). Our observations from the various cell growth experiments suggested that loss of ACSM1 and ACSM3 caused cytotoxicity, which we tested using Annexin V/7-AAD assays. Annexin V binds to phosphatidylserine on the surface of cells that are undergoing multiple types of programmed cell death, including apoptosis, necroptosis and ferroptosis^25^. Targeting of either factor caused a significant increase in cell death in multiple cell line models of PCa (Figure 3F). Collectively, these findings reveal that ACSM1 and ACSM3 are required for growth and survival of PCa cells.

**Figure 3.**
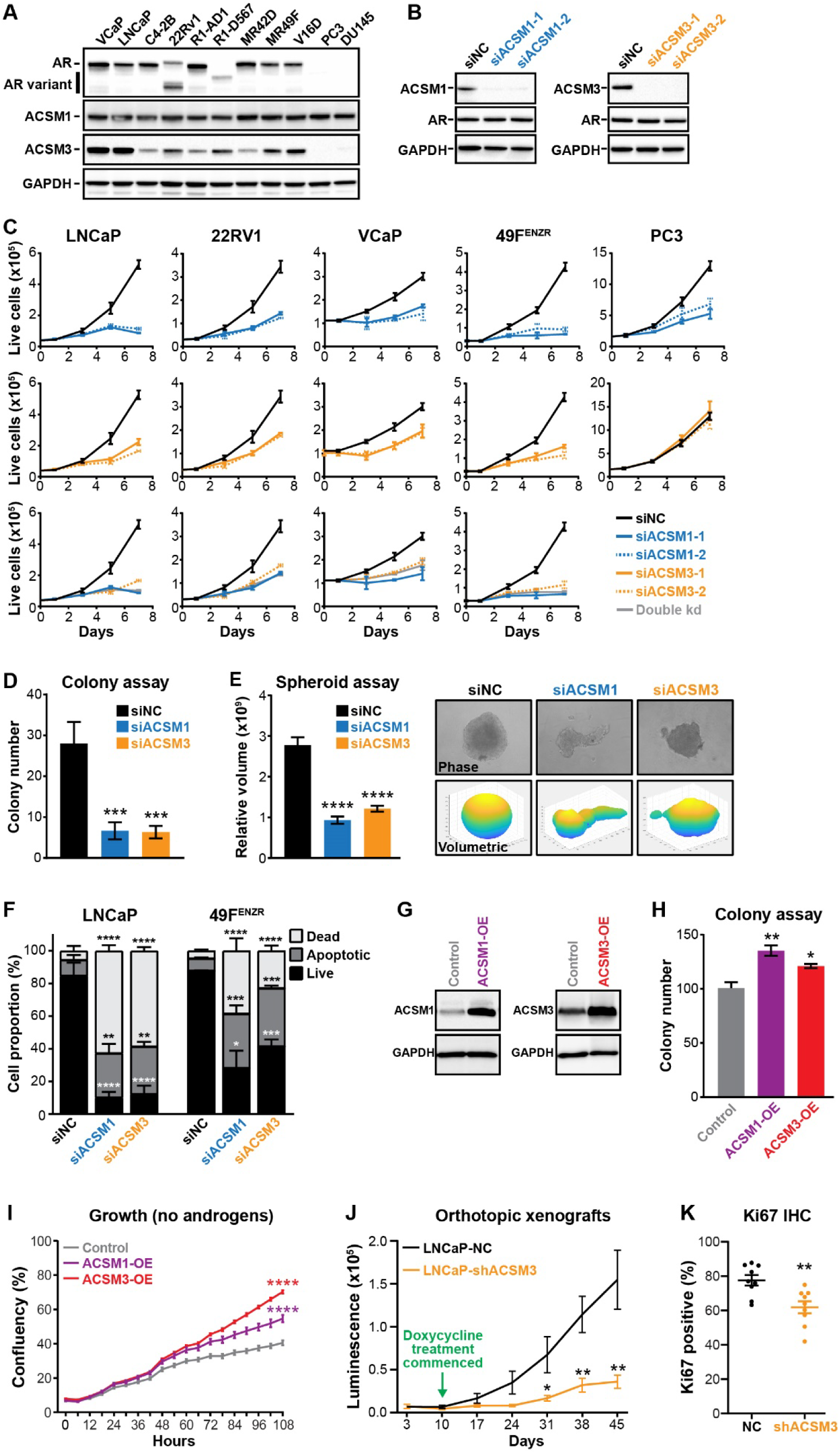
ACSM1 and ACSM3 are pro-growth and pro-survival factors in prostate cancer. **(A)** Western blotting of AR, ACSM1 and ACSM3 protein expression in PCa cell lines. GAPDH was used as a loading control. **(B)** Efficacy of siRNAs targeting ACSM1 and ACSM3. 10 nM of two distinct siRNAs per gene (siACSM1-1, siACSM1-2, siACSM3-1 and siACSM3-2) were transfected into LNCaP cells for 72 h, after which ACSM1 and ACSM3 protein levels were evaluated by Western blotting. GAPDH was used as a loading control. **(C)** Loss of ACSM1 and ACSM3 inhibits prostate cancer growth, as evaluated by Trypan blue assays. LNCaP, 22Rv1, VCaP, 49F^ENZR^ and PC3 cells were transfected with 2 distinct siRNAs per gene and Trypan blue growth assays were performed. Error bars represent SEM. Unpaired t tests were used to compare growth of siACSM1/siACSM3 relative to siNC at day 7. **(D)** Loss of ACSM1 and ACSM3 inhibits LNCaP colony formation. Cells were grown for 2 weeks, washed with PBS, fixed with paraformaldehyde and stained with 1% crystal violet for 30 min. Colonies were counted manually. Data shown is representative of 3 independent experiments. Unpaired t tests were used to compare colony formation of siACSM1/siACSM3 relative to siNC. **(E)** Loss of ACSM1 and ACSM3 inhibits growth of LNCaP spheroids. Spheroid volumes were determined using the ReViSP software. Data shown on left is representative of 3 independent experiments. Representative image of spheres and volumetric analyses are shown in right. Unpaired t tests were used to compare spheroid volume of siACSM1/siACSM3 relative to siNC. **(F)** ACSM1 and ACSM3 knockdown causes apoptosis of LNCaP cells, as determined using flow cytometry-based Annexin V/7-AAD assays. Data represents the mean ± SEM of triplicate samples and are representative of 3 independent experiments. Dead cell proportions were compared to vehicle using ANOVA and Dunnett’s multiple comparison tests. **(G)** Western blotting showing over-expression of ACSM1 and ACSM3 in stably transduced LNCaP cells. GAPDH was used as a loading control. **(H)** Over-expression of ACSM1 and ACSM3 promotes colony formation. Colonies (grown over 2 weeks) were visualised and counted as in (D). Colony numbers were compared to control cells using ANOVA and Dunnett’s multiple comparison tests. **(I)** Over-expression of ACSM1 and ACSM3 promotes growth of LNCaP cells in androgen-depleted media. Confluency over a period of 108 hours was measured using an Incucyte live-cell imaging and analysis platform. Confluency was compared to control cells (at 108 hours) using ANOVA and Dunnett’s multiple comparison tests. **(J)** Knockdown of ACSM3 inhibits growth of LNCaP intra-prostatic xenografts. The graph represents luciferase intensity as assessed by whole-animal bioluminescent imaging over time in mice injected with shNC (control) cells (n = 8) or shACSM3 cells (n = 10). Data are presented as mean ± SEM. The shACSM3 and shNC groups were compared at indicated time-points using unpaired t tests. **(K)** Quantitation of immunostaining for the proliferative marker Ki67 in LNCaP-NC and LNCaP-shACSM3 xenografts. The shACSM3 and shNC groups were compared at day 45 using an unpaired t test. For all statistical tests: *, p < 0.05; **, p < 0.01; ***, p < 0.001; ****, p < 0.0001.

To model increased activity of ACSM1/3, a common feature of PCa (Figure 2), we generated LNCaP cells stably modified to express higher levels of either factor (Figure 3G). In contrast to the findings from the loss of function experiments, ACSM1 and ACSM3 over-expressing cells (ACSM1-OE and ACSM3-OE) exhibited a slightly elevated growth rate and an augmented ability to form colonies (Figure 3H, Supplementary Figure S2A). Enhanced proliferative capacity of these cells was particularly evident in androgen-depleted conditions (Figure 3I), suggesting that elevated ACSM1 or ACSM3 in patient tumours could influence response to ADT.

Encouraged by the *in vitro* data across a diverse range of PCa models, we investigated the requirement of ACSM1/3 for prostate cancer tumour growth *in vivo*. Luciferase-tagged LNCaP cells were engineered with lentiviruses to express doxycycline-inducible shRNAs specific for ACSM1 or ACSM3. LNCaP-shACSM3 cells exhibited robust knockdown of ACSM3 in response to doxycycline, and this shRNA-mediated knockdown recapitulated the effects on growth and colony forming ability elicited by siRNAs (Supplementary Figure S2B-D). By contrast, LNCaP-shACSM1 cells did not exhibit robust down-regulation of ACSM1 (Supplementary Figure S2B), and hence we did not proceed to *in vivo* experiments with this model. LNCaP-shACSM3 cells were injected directly into the prostate and the growth of orthotopic xenografts was measured by monitoring of luciferase expression. After a period of 10 days to establish the xenografts, mice were fed doxycycline to induce knockdown of ACSM3. Loss of ACSM3 led to a striking decrease in the growth of LNCaP tumour xenografts over a period of 48 days (Figure 3J), and this was associated with decreased levels of the proliferative marker Ki67 in tumours (Figure 3K; Supplementary Figure S2E).

### ACSM1 and ACSM3 are required for energy production and control lipid composition in prostate cancer cells

Members of the ACSM family catalyse the activation of MCFAs to produce acyl-CoA, a reaction that is indispensable in fatty acid utilisation^26^. However, the relevance of MCFAs to the growth of PCa cells is poorly understood. Multiple studies have shown that dietary MCFAs can inhibit the growth of cancer cells^27, 28^, whereas a study of prostate epithelial lines found that medium chain triglycerides enhance the growth of benign cells but have little effect on cancer cells^29^. We found that moderate doses of the MCFAs sodium octanoate and decanoic acid (C10:0, also known as capric acid) could weakly stimulate growth of PCa cells, whereas high doses were growth inhibitory and caused cell death (Supplementary Figure S3A-C).

Having established that MCFAs can influence PCa proliferation, we undertook a series of experiments to interrogate bioenergetic metabolic alterations in response to modulation of ACSM1 or ACSM3. Using a luminescence-based assay, we found that knockdown of either ACSM1 or ACSM3 resulted in a decrease in total cellular ATP (Supplementary Figure S3C). Next, we used extracellular flux analysis of live cells to determine how ACSM1/3 influence oxidative phosphorylation (OXPHOS) in live cells. In substrate-limiting conditions, addition of sodium octanoate (NaOc) to a range of PCa cell line models (49F^ENZR^, 22Rv1 and LNCaP) substantially increased the oxygen consumption rate (OCR), most notably the maximal respiration rate, demonstrating that this lipid can be used as a substrate for OXPHOS (Figure 4A-C). Importantly, knockdown of ACSM1 or ACSM3 blunted the ability of cells to metabolise sodium octanoate (Figure 4A-B). Conversely, over-expression of ACSM1 or ACSM3 in LNCaP cells enhanced respiration in response to sodium octanoate (Figure 4C). Collectively, these experiments demonstrate the importance of ACSM1/3 in the metabolism of medium chain lipids.

**Figure 4.**
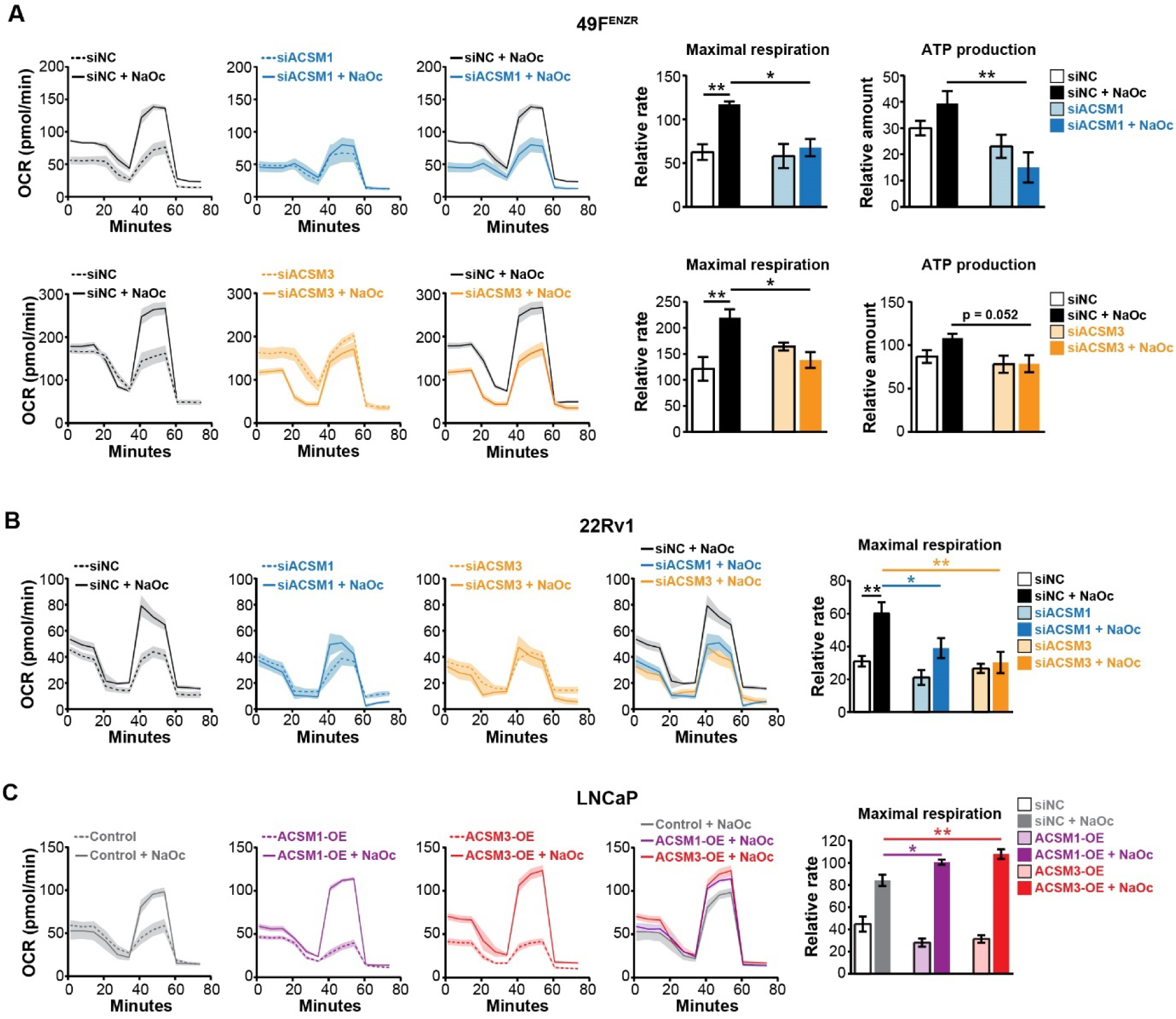
ACSM1 and ACSM3 enhance mitochondrial oxidative phosphorylation. **(A)** Loss of ACSM1 or ACSM3 reduces oxygen consumption rate (OCR), maximal respiration and ATP production in 49F^ENZR^ cells. Cells were transfected with siACSM1, siACSM3 or siNC for 72 hr, then starved in substrate limited medium for 24 hr; the assay was run in fatty acid oxidation assay medium supplemented with sodium octanoate (NaOc). OCR was normalised to cell number. **(B)** Loss of ACSM1 or ACSM3 reduces oxygen consumption rate (OCR), maximal respiration and ATP production in 22Rv1 cells. The assay was performed as described in (A). **(C)** Over-expression of ACSM1 or ACSM3 increases oxygen consumption rate (OCR), maximal respiration and ATP production in LNCaP cells. Data in bar graphs are mean ± SEM. One way ANOVA and Tukey’s multiple comparisons test were used to compare groups. For all statistical tests: *, p < 0.05; **, p < 0.01.

Given their purported functions in metabolism of medium chain fatty acids and thereby precursor availability for lipid synthesis, the influence of ACSM1 and ACSM3 on levels and composition of lipids in PCa cells was evaluated. First, LNCaP cells were stained with BODIPY 493/503 to measure total neutral lipids, such as triglycerides and cholesteryl esters. Over-expression of ACSM1/3 caused a decrease in cellular neutral lipids, presumably due to enhanced capacity for lipid activation and utilisation (Figure 5A), whereas knockdown of either factor caused cells to accumulate lipids, indicative of defective lipid metabolism (Figure 5A). We then used a quantitative mass spectrometric approach to profile the PCa lipidome in more detail. Paralleling the BODIPY staining, knockdown or over-expression of ACSM1/3 increased or decreased, respectively, the global lipid content of PCa cells (Figure 5B; Supplementary Figure S4; Dataset S1). Grouping of lipids by saturation status revealed that modulation of either enzyme affected saturated FAs, MUFAs and PUFAs (Figure 5C; Dataset S1). Almost all lipid classes accumulated in response to ACSM1/3 knockdown and decreased with ACSM1/3 over-expression (Figure 5D; Supplementary Figure S4A-B; Dataset S1). Although there was significant overlap in lipids altered following modulation of ACSM1 and ACSM3, unique changes were observed (Supplementary Figure S4C-D). A set of lipids that were concordantly altered in all 4 conditions (i.e. elevated in response to loss of ACSM1/3 and decreased in response to over-expression of ACSM1/3) was enriched for various sphingomyelin (SM) species (Supplementary Figure S4C-D), which play important roles in cell membranes^30^. Overall, these data demonstrate that the degree of ACSM1/3 expression and activity is an important regulator of the PCa cellular lipidome and that there are both shared and non-redundant effects of each enzyme in relation to specific lipid species. Importantly, increased expression of ACSM1/3 endows PCa cells with an augmented capacity to utilise fatty acids for energy and other purposes, the process of which prevents accumulation of cellular lipids to toxic levels.

**Figure 5.**
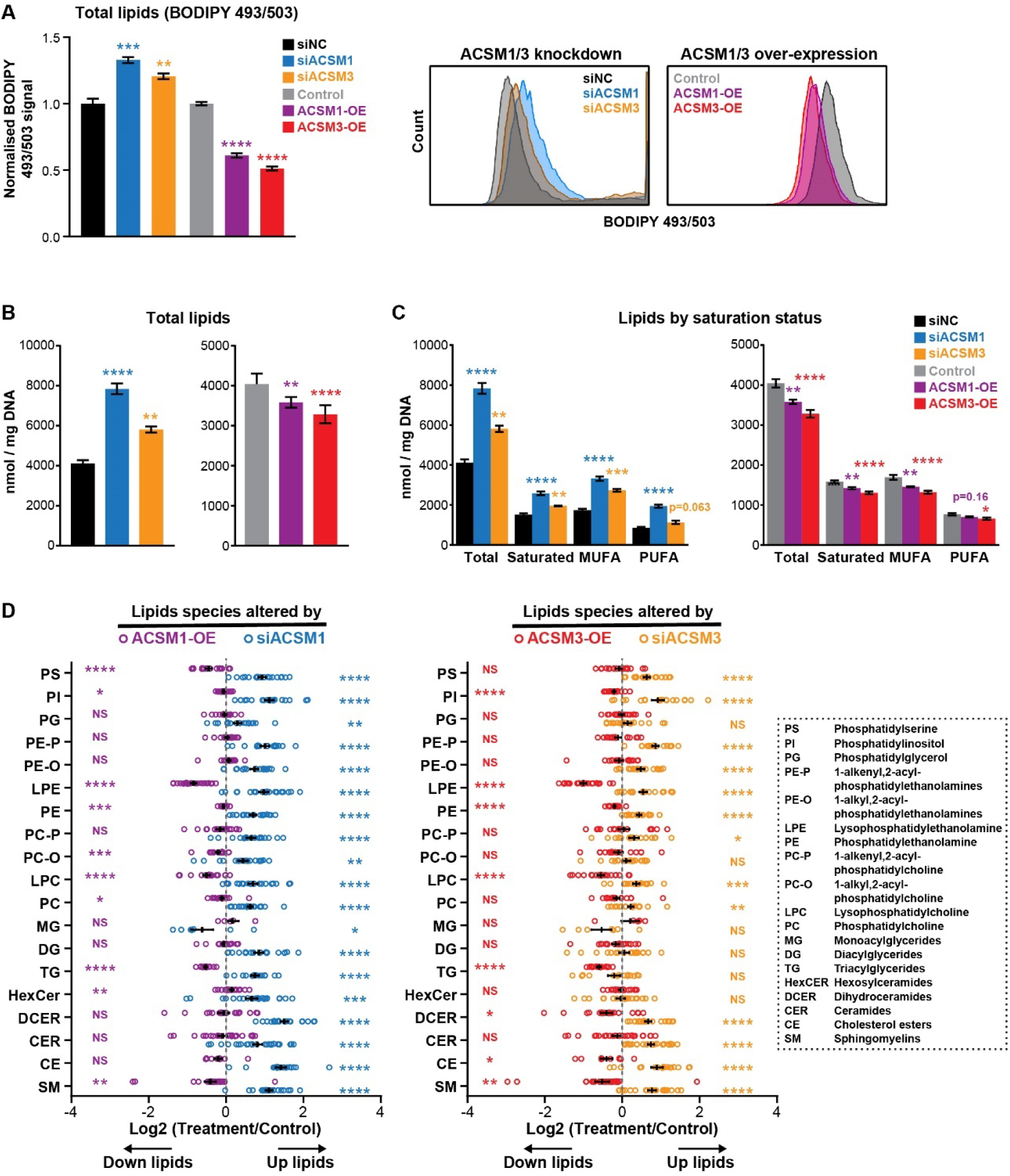
ACSM1 and ACSM3 are major regulators of the prostate cancer lipidome. **(A)** Neutral lipid content of LNCaP cells in response to ACSM1/3 knockdown (siACSM1, siACSM3) or over-expression (ACSM1-OE and ACSM3-OE) was detected using BODIPY 493/503 dye and flow cytometry. One way ANOVA and Tukey’s multiple comparisons test were used to compare groups. Flow cytometric graphs are shown on right. **(B)** Abundance of total lipids in response to knockdown or over-expression of ACSM1 and ACSM3, as determined by mass spectrometry. Data in bar graphs are mean ± SEM. Unpaired t tests were used to compare siACSM1/siACSM3 relative to siNC. **(C)** Abundance of lipids by saturation status. Data in bar graphs are mean ± SEM. Unpaired t tests were used to compare siACSM1/siACSM3 relative to siNC or ACSM1-OE/ACSM3-OE relative to control. **(D)** Abundance of lipids by class. Data is expressed as Log2 of treatment/control for each lipid species, which are indicated by circles. Black lines are mean ± SEM. One-sample t-tests were used to determine significant changes in abundance for each class.

As another strategy to evaluate the role of ACSM1/3 in PCa cell metabolism, we undertook metabolomic profiling using mass spectrometry. Knockdown of either ACSM1 or ACSM3 caused accumulation of free CoA, consistent with the purported function of ACSM enzymes in generating fatty acyl-CoaA esters (Supplementary Figure S5A). Cells lacking ACSM1/3 also exhibited increased levels of the glucose-6-phosphate, fructose-1,6-bisphosphate and intracellular lactate, which are intermediates and the end-product of glycolysis, and amino acids (Supplementary Figure S5B-C). These latter observations are consistent with the idea that loss of ACSM1/3 causes PCa cells to undergo a switch from OXPHOS to glycolysis, a reported consequence of a reduced ability to utilise FAO for energy production^31, 32^.

### Targeting of ACSM1 and ACSM3 induces lipid peroxidation and ferroptosis

Our metabolic and lipidomic approaches indicated that targeting ACSM1 and ACSM3 causes remodelling of PCa metabolism and the lipidome, both of which could induce oxidative stress. In support of this concept, loss of ACSM1/ACSM3 activity dramatically increased the levels of reactive oxygen species (ROS), as determined by a flow cytometry-based assay, in multiple models of PCa (Figure 6A-B). Mitochondrial superoxide, measured using MitoSOX assays, was a significant component of the increased cellular ROS (Figure 6C). Increased ROS was consistent with a substantial decrease in the antioxidant glutathione (Figure 6D) and an increase in the NAD+:NADH ratio (Figure 6E). Interestingly, the depletion of glutathione could not be explained by the availability of its constituent amino acids (cysteine, glutamate and glycine; Figure 6F). Collectively, these data provide evidence for mitochondrial dysfunction in response to knockdown of ACSM1/ACSM3 and reveal that both enzymes are required to maintain cellular redox homeostasis.

**Figure 6.**
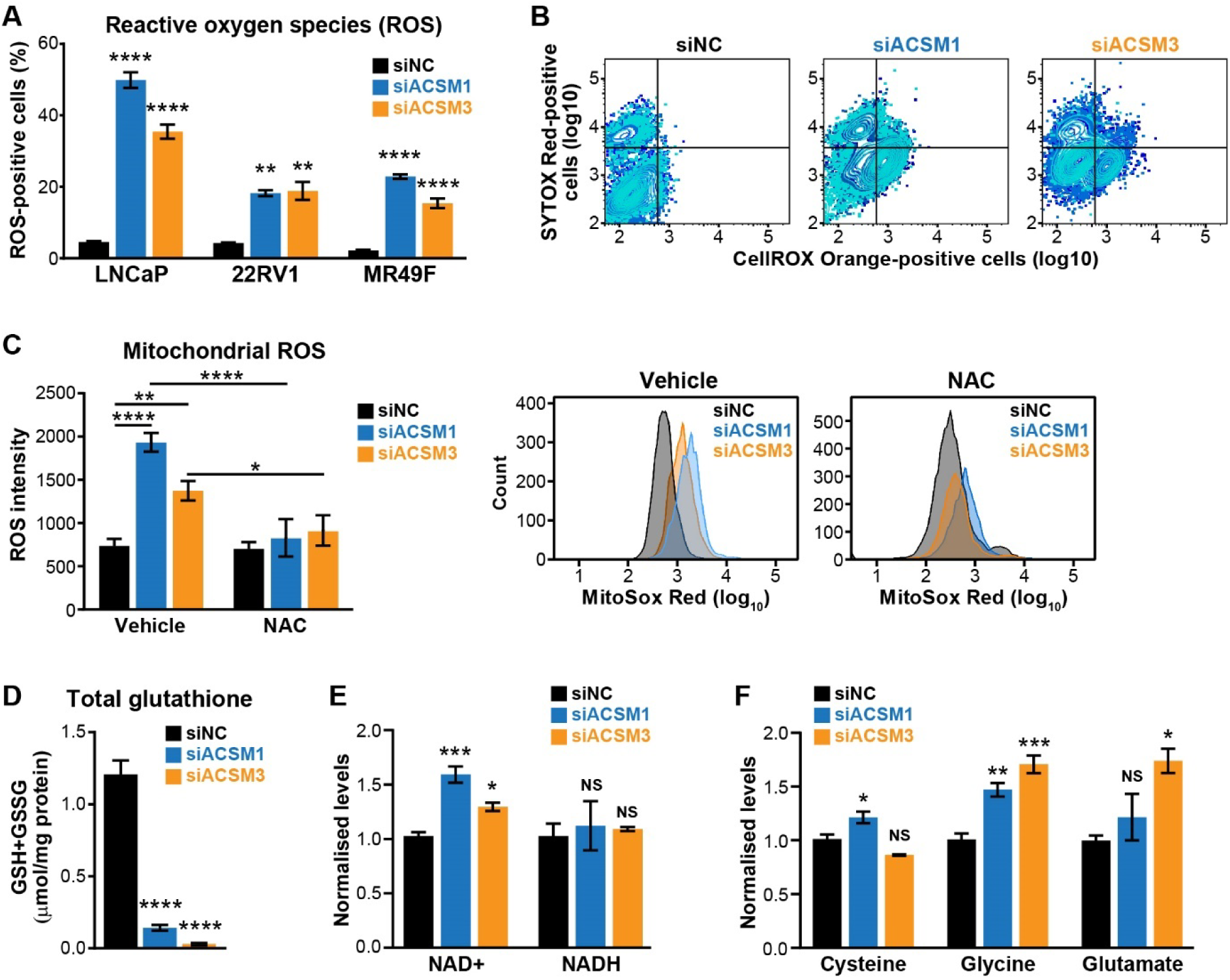
ACSM1 and ACSM3 maintain redox homeostasis in prostate cancer cells. **(A-B)** Loss of ACSM1 and ACSM3 causes accumulation of ROS (A), as determined by a CellROX orange flow cytometric assay. Representative flow cytometry data for LNCaP cells are shown in (B). Data shown in (A) are mean ± SEM. Unpaired t tests were used to compare siACSM1/siACSM3 relative to siNC. **(C)** Loss of ACSM1 and ACSM3 causes accumulation of mitochondrial ROS (left), evaluated using a MitoSOX assay to measure mitochondrial superoxide levels. Representative flow cytometry data are shown on left. One way ANOVA and Tukey’s multiple comparisons test were used to compare groups. **(D)** Loss of ACSM1 and ACSM3 causes a reduction in total glutathione (reduced glutathione (GSH) + oxidized glutathione (GSSG). Data are mean ± SEM. Unpaired t tests were used to compare siACSM1/siACSM3 relative to siNC. **(E)** Loss of ACSM1 and ACSM3 alters levels of NAD+, measured using GC QQQ targeted metabolomics. Data are mean ± SEM. Unpaired t tests were used to compare siACSM1/siACSM3 relative to siNC. **(F)** Levels of amino acid constituents of glutathione, measured using GC QQQ targeted metabolomics. Data are mean ± SEM. Unpaired t tests were used to compare siACSM1/siACSM3 relative to siNC. For all statistical tests: *, p < 0.05; **, p < 0.01; ***, p < 0.001; ****, p < 0.0001.

The combination of increased ROS and accumulation of lipids, in particular those containing PUFAs, in response to depletion of ACSM1/3 is likely to result in lipid peroxidation. To test this concept, we measured a marker of lipid peroxidation, malondialdehyde (MDA)^33^, and found that it was strongly induced in response to knockdown of ACSM1 and ACSM3 (Figure 7A-B). This observation was confirmed by staining cells with BODIPY-C11, which binds to phospholipid hydroperoxides in cell membranes (Figure 7C). Lipid peroxidation is a hallmark of ferroptosis, an iron-dependent, non-apoptotic form of cell death^34^. Treatment of cells with two inhibitors of ferroptosis, ferrostatin-1 and N-acetylcysteine (NAC), reversed the effect of siRNA treatment on cell death and ROS production in multiple models of PCa (Figure 7D; Supplementary Figure S6), confirming that ablation of ACSM1/3 leads to ferroptotic cell death.

**Figure 7.**
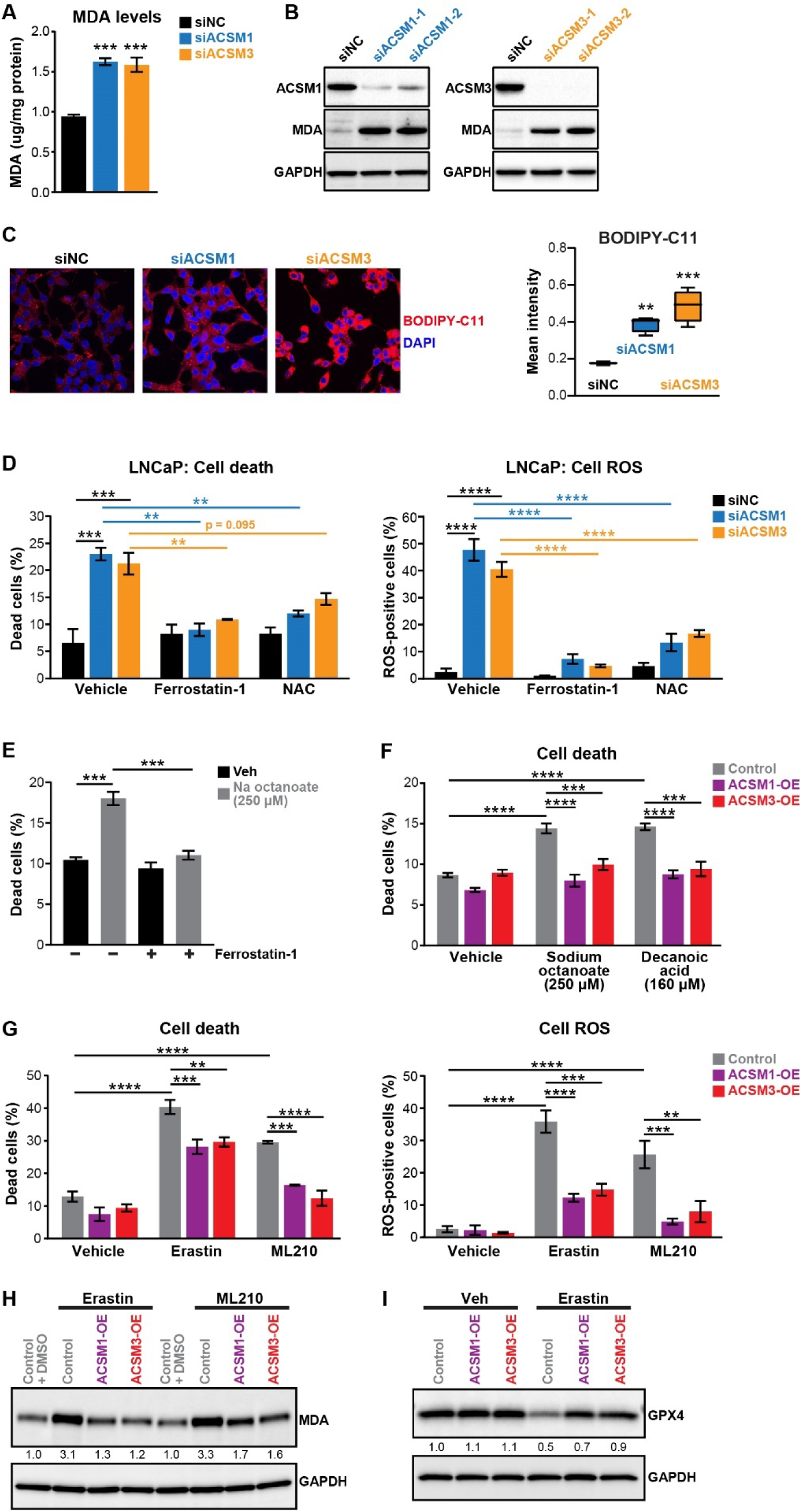
ACSM1 and ACSM3 protect prostate cancer cells from ferroptosis. **(A-B)** Loss of ACSM1 and ACSM3 causes accumulation of malondialdehyde (MDA), as measured using a colourimetric assay (A) and Western blotting (B). For (A), data are mean ± SEM and unpaired t tests were used to compare siACSM1/siACSM3 relative to siNC. For (B), GAPDH was used as a loading control. **(C)** Loss of ACSM1 and ACSM3 causes accumulation of membrane phospholipid hydroperoxides, as evaluated using BODIPY-C11 immunofluorescence in LNCaP cells. Quantitation of BODIPY-C11 signal is shown on right. Boxes represent 25^th^-75^th^ percentiles, the line within boxes is the median, and top and bottom whiskers are maximum and minimum values, respectively. One way ANOVA and Tukey’s multiple comparisons test were used to compare groups. **(D)** Ferrostatin-1 (1.25 µM) and NAC (5 µM) rescue cell death and ROS induction mediated by loss of ACSM1 and ACSM3 in LNCaP cells. Cell death and ROS were measured by flow cytometric assessment of SYTOX Red- and CellROX-orange stained cells, respectively. Data are mean ± SEM. One way ANOVA and Tukey’s multiple comparisons test were used to compare groups. **(E)** Effect of sodium octanoate (Na Octanoate) on LNCaP cell death, as measured by flow cytometric staining of 7-AAD. Data are mean ± SEM. One way ANOVA and Tukey’s multiple comparisons test were used to compare groups. **(F)** Rescue of cell death elicited by fatty acids by over-expression of ACSM1 and ACSM3. Death was measured as in F. Data are mean ± SEM. One way ANOVA and Tukey’s multiple comparisons test were used to compare groups. **(G)** Over-expression of ACSM1 and ACSM3 rescues cells from cell death and ROS induced by ferroptosis-inducing compounds. Cells were treated with 10 µM Erastin or 2.5 µM ML210 for 3 days prior to measurement of cell death or ROS using flow cytrometry. Data are mean ± SEM. One way ANOVA and Tukey’s multiple comparisons test were used to compare groups. **(H)** Over-expression of ACSM1 and ACSM3 reverses elevation of MDA levels caused by ferroptosis-inducing compounds. Cells were treated with 10 µM Erastin or 2.5 µM ML210 for 4 days prior to Western blotting. GAPDH was used as a loading control. **(G)** Over-expression of ACSM1 and ACSM3 maintains GPX4 in the presence of ferroptosis-inducing compounds. Cells were treated with 10 µM Erastin for 4 days prior to Western blotting. GAPDH was used as a loading control. For all statistical tests: *, p < 0.05; **, p < 0.01; ***, p < 0.001; ****, p < 0.0001.

Our earlier findings suggested that media supplemented with high levels of MCFAs is toxic to PCa cells (Supplementary Figure S3), a finding that recapitulates recent work in glioblastoma^35^. Considering these observations in combination with the ability of ACSM1/3 to enhance cell survival, we speculated that MCFA lipotoxicity may, at least in part, be a result of ferroptosis. Supporting this hypothesis, cell death elicited by sodium octanoate and decanoic acid was partially rescued by ferrostatin-1 (Figure 7E). Over-expression of ACSM1/3 ameliorated MCFA lipotoxicity (Figure 7F), indicating that one pro-survival function of these lipid metabolic enzymes is to prevent accumulation of C8/C10 lipids to levels that stimulated ROS production, lipid peroxidation and thereby elicit ferroptosis.

To further model how increased ACSM1/3 expression in PCa cells influences ferroptosis, we evaluated the response of our stable over-expression cell lines to two distinct ferroptosis-inducing drugs: Erastin inhibits the cystine-glutamate antiporter system (via VDAC2 and VDAC3) whereas ML210 inhibits glutathione peroxidase 4 (GPX4), a selenoenzyme that uses reduced glutathione to eliminate lipid peroxides and thereby acts as a guardian against ferroptosis^36, 37^. Cells over-expressing ACSM1/3 were significantly less susceptible to these compounds, exhibiting resistance to cell death and ROS production (Figure 7G) and reduced levels of the lipid peroxidation marker MDA (Figure 7H). Treatment with ferroptosis-inducing compounds is known to decrease expression of GPX4, including in PCa cells^38^. Interestingly, over-expression of ACSM1/3 prevented Erastin-mediated down-regulation of GPX4 (Figure 7I). Collectively, these orthogonal and complementary approaches demonstrate that a key function of ACSM1/3 is to ameliorate lipid peroxidation and thereby protect PCa cells against ferroptosis.

### ACSM1 and ACSM3 protect prostate cancer cells from redox stress and lipid peroxidation induced by AR inhibition

A major role of the AR signalling axis in highly metabolically active PCa cells is to protect against oxidative stress^39^. Indeed, inhibiting AR signalling enhances lipid peroxidation^40^, a phenomenon we also observed by measuring BODIPY-C11 and MDA in LNCaP cells treated with the AR antagonists enzalutamide or darolutamide (Figure 8A-B). Importantly, over-expression of ACSM1 or ACSM3 alleviated lipid peroxidation (Figure 8C-D) and iron-dependent accumulation of ROS (Figure 8E) elicited by enzalutamide or darolutamide, revealing a potential role of these enzymes in enabling therapy-resistant tumour phenotypes.

**Figure 8.**
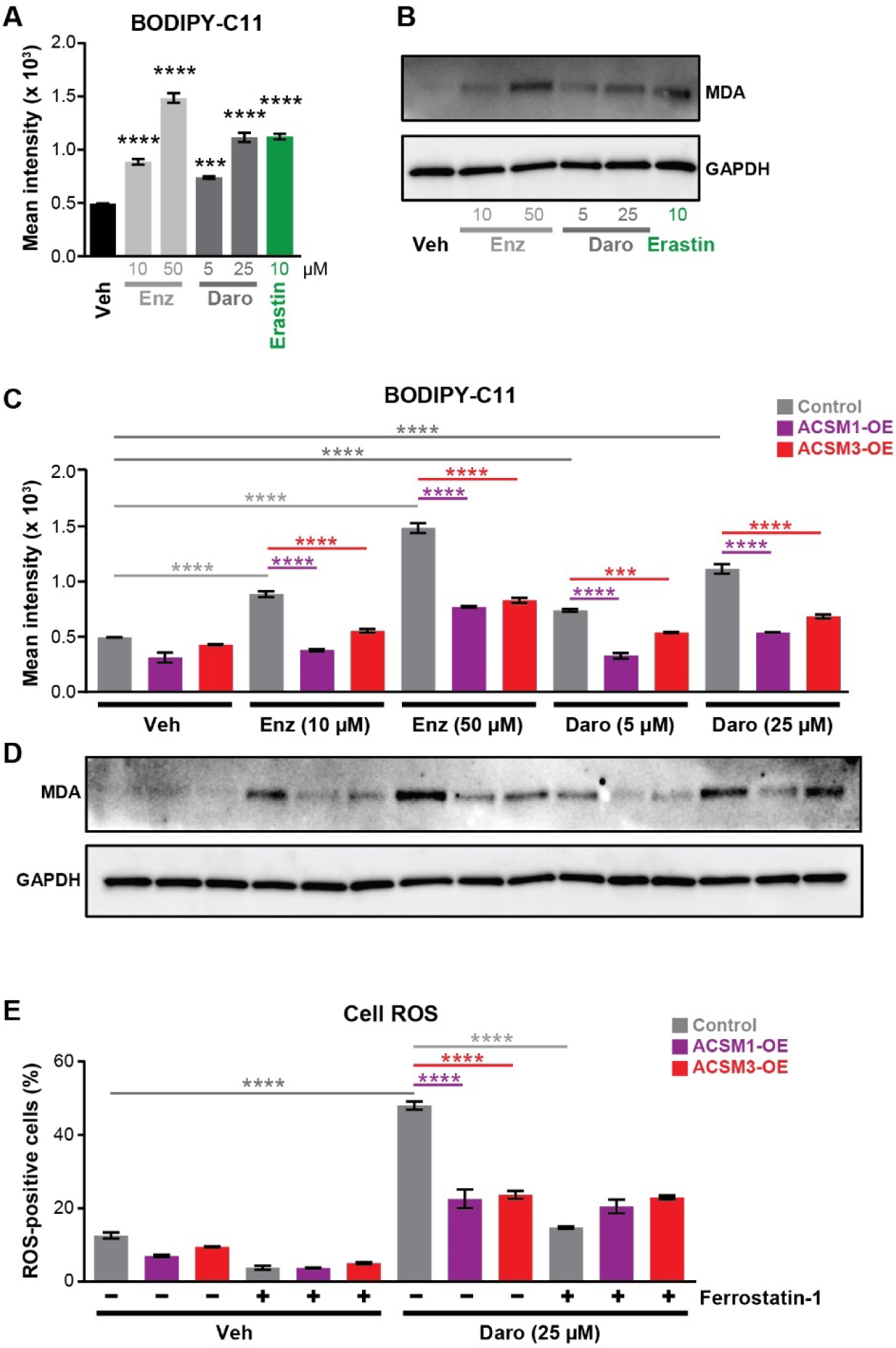
ACSM1 and ACSM3 protect prostate cancer cells from redox stress and lipid peroxidation induced by AR inhibition. **(A)** AR antagonists cause accumulation of membrane phospholipid hydroperoxides. Cells were treated with AR antagonists (enzalutamide, Enz or darolutamide, Daro) or Erastin for 4 days prior to measuring BODIPY-C11 levels by flow cytometry. Data are mean ± SEM. One way ANOVA and Tukey’s multiple comparisons test were used to compare treatments to vehicle. **(B)** AR antagonists cause accumulation of malondialdehyde (MDA). Cells were treated as in A prior to evaluation of protein levels by Western blotting. GAPDH was used as a loading control. **(C-D)** Over-expression of ACSM1 or ACSM3 reverses lipid peroxidation (C) and MDA accumulation (D). Experiments were performed as in A-B. **(E)** Over-expression of ACSM1 and ACSM3 rescues cells from ROS accumulation elicited by darolutamide (Daro). Cells were treated with darolutamide for 4 days, in the presence or absence of ferrostatin-1, prior to measuring ROS using flow cytrometry. Data are mean ± SEM. One way ANOVA and Tukey’s multiple comparisons test were used to compare groups. For all statistical tests: *, p < 0.05; **, p < 0.01; ***, p < 0.001; ****, p < 0.0001.

## DISCUSSION

The AR signalling axis is a major regulator of lipid metabolic processes in prostate cancer, but the molecular mechanisms underlying this regulation and its relevance to disease progression remain poorly understood. In this study, we demonstrated that AR directly regulates the levels of two mitochondrial acyl-CoA synthetases, ACSM1 and ACSM3, and found that these enzymes play a critical role in lipid metabolism and protecting cells against oxidative stress, lipid peroxidation and ferroptosis induced by MCFAs, ferroptosis-inducing drugs and AR-targeted therapies. These findings are significant because they provide novel insights into the interplay between lipid metabolism and stress responses in prostate cancer as well as revealing two new targets for future therapies.

The functions of ACSM1 and ACSM3 in normal physiology are poorly understood. Indeed, the ACSM family as a whole is highly understudied compared to most other metabolic enzymes^41^. The *ACSM1* and *ACSM3* genes share a high degree of homology and structural similarity and are co-located on chromosome 16p13.1, suggesting that they arose from a common ancestor by gene duplication^42^. Although early *in vitro* studies demonstrated that human ACSM1 prefers hexanoate (6 carbons) as a substrate^43^ whereas the mouse orthologues of ACSM1 and ACSM3 prefer octanoate (8 carbons) and isobutyrate (4 carbons) as substrates, respectively^42^, it has become apparent that there is significant chain length diversity and overlap in ACSM substrates^44, 45^. In terms of pathophysiological functions, a major research focus has been establishing and understanding the association between *ACSM1* and *ACSM3* gene polymorphisms and components of the metabolic syndrome (i.e. hypertension, hypertriglyceridemia, hypercholesterolemia and obesity)^46–49^. Very little is known about the relevance of ACSMs in cancer. ACSM1 has been reported to be a marker of PCa^50, 51^ and apocrine breast cancer^52^, whereas ACSM3 has been implicated as a tumour suppressor in hepatocellular carcinoma based on its down-regulation in this disease context^53, 54^.

Ours is the first study to undertake integrative functional characterisation of ACSM1 and ACSM3 in any cancer type. We found that both enzymes are encoded by AR-regulated genes, which may explain the observation that prostate tumours exhibited the highest ACSM1/3 mRNA levels in our pan-cancer survey. AR stimulates the transcription of genes encoding many different factors involved in distinct aspects of lipid metabolism, including lipid synthesis, elongation, uptake and trafficking^55^. A recent study identified long chain acyl-CoA synthetases as members of the androgen signalling axis; more specifically, ACSL4 was repressed whereas ACSL3 was activated by AR activity^56^. This work, alongside our new research, reveals AR regulation of acyl-CoA synthetases to be a previously unappreciated and important mechanism by which androgen signalling influences lipid metabolism in PCa cells.

One important function of ACSM1/3 revealed herein relates to energy production via FAO. We found that PCa cells can utilise octanoate as an energy source, a finding that may have significant relevance to patients since these lipid species circulate at significant levels in serum and their uptake is not subject to negative feedback regulation^35, 57^. The ability of ACSM1/3 to enhance FAO is a straightforward explanation for their pro-growth activity and aligns with the observation that PCa cells exhibit substantially increased rates of FAO compared to non-malignant prostate cells^58, 59^. Importantly, the concept that MCFAs can fuel tumour growth is increasingly well recognised; for example, a recent study found that FAO of MCFAs serves as a crucial energy source for glioblastoma^35^. Thus, enzymes that can enhance oncogenic MCFA metabolism, such as ACSM1/3 (this study) and medium-chain acyl-CoA dehydrogenase^35^, warrant further investigation as therapeutic targets in oncology, particularly for tumours characterised by lipid-rich environments such as the prostate.

In recent years, ferroptosis has emerged as a crucial non-apoptotic mode of cell death^60^. A hallmark of ferroptosis is peroxidation of lipids, in particular PUFA phospholipids in cell membranes, in a manner dependent on accumulation of iron^60^. Using both loss- and gain-of-function experiments in combination with compounds that can activate or suppress ferroptosis, our study revealed that ACSM1/3 are potent suppressors of ferroptotic cell death in PCa. Mechanistically, our findings demonstrate that ACSM1/3 have pleiotropic protective effects against PCa ferroptosis. First, these enzymes constrain accumulation of ROS, which are required for lipid peroxidation, and assist in maintaining antioxidants such as reduced glutathione and NADH. Glutathione plays a particularly critical role in protecting cells against ferroptosis since it is used by GPX4 to reduce lipid peroxides to non-toxic lipid alcohols^61^. We propose that ACSM1/3 stimulates mitochondrial function and the tricarboxylic acid cycle, resulting in generation of NADH that can be used to reduce oxidized glutathione. Supporting this concept, mitochondrial dysfunction is linked to reduced glutathione levels in a variety of disorders and systems^62^. Second, our study found that ACSM1/3 are required to maintain normal levels of GPX4, which plays a critical role in eliminating lipid peroxides. It has been proposed that increased OXPHOS activity is met with an increased need for GPX4^63, 64^, which may explain the positive link between ACSM1/3 activity and this key anti-ferroptotic factor. Third, by promoting turnover of MCFAs, ACSM1/3 would suppress lipotoxicity caused by accumulation of these lipid species. Indeed, we found that high levels of MCFAs inhibit growth and cause death of prostate cancer cells, at least partly by eliciting ferroptosis. This is reminiscent of toxicity associated with C10 and C12 lipid accumulation in glioblastoma cells, although no evidence of ferroptosis was observed in that disease context^35^. More broadly, we propose that the functions of ACSM1/3 in lipid metabolism and fatty acid oxidation, redox homeostasis and ferroptosis are likely to be highly interconnected and that dissecting these links will yield new insights into the unique pathometabolic features of PCa cells.

Our study also provided new insight into the mode of action of AR antagonists, demonstrating that they can cause lipid peroxidation in PCa cells. Down-regulation of ACSM1/3 in response to inhibition of AR transcriptional activity may represent one mechanism underlying this phenomenon, underscored by the observation that ACSM1/3 over-expression can mitigate lipid peroxidation and redox stress caused by enzalutamide and darolutamide. These findings are also consistent with recent work by Stoyanova and colleagues, who demonstrated that combining ferroptosis-inducing drugs with enzalutamide exhibited significantly greater anti-tumour activity *in vivo* compared to the single agents^65^. More broadly, these observations highlight the potential utility of targeting ferroptosis in the castration-resistant disease context. Indeed, strategies that induce ferroptosis may be particularly useful in CRPC since: i) therapy-resistant phenotypes, in multiple cancer types and in response to diverse therapies, have been associated with increased sensitivity to ferroptosis due to altered fatty acid membrane composition^2, 66–68^; and ii) acquisition of resistance to AR-targeted therapies has been linked to increased uptake and mobilisation of PUFAs^40^ as well as elevated lipid catabolism more broadly^69^.

The findings described herein further reinforce the emerging concept that PCa cells must acquire strategies to overcome ferroptotic vulnerability whilst maintaining oncogenic lipid metabolism. Beyond selection for high expression and activity of ACSM1/3, recent reports demonstrated that another enzyme required for fatty acid oxidation, DECR1, serves to protect PCa cells from ferroptosis^31, 70^. In contrast to ACSM1/3, DECR1 is an AR-repressed factor, although (paralleling ACSM1/3) it is upregulated during PCa development and progression and serves to process PUFAs, key substrates of lipid peroxidation during ferroptosis, for catabolism. Another adaptive survival response against ferroptosis in PCa cells is regulation of arachidonic acid metabolism and increased production of the anti-ferroptotic factor prostaglandin E2 by ATF6α and PLA2G4A^71^. Collectively, the identification of distinct ferroptosis-inhibiting lipid metabolic pathways in PCa exemplifies metabolic evolution in malignancy as well as revealing novel pathways that could be therapeutically targeted.

A limitation of our study was the failure to detect medium chain FAs or their activated forms by mass spectrometry, which makes it difficult to pinpoint specific substrates of ACSM1/3. However, reports suggest that there is significant overlap in the chain length specificity of ACSMs and ACSLs^44, 45^; with this in mind and based on our lipidomic data, we speculate that ACSM1/3 directly catalyze activation of both medium and long chain FAs in PCa cells. Further examination of the enzymatic activities of ACSM1/3, including identification of shared and unique substrates, will be important step to fully understand their roles in PCa lipid metabolism.

In summary, our study defines the first direct link between the AR signalling axis and enzymes involved in the activation of medium-chain fatty acids. Importantly, the dual roles of ACSM1 and ACSM3 in both energy production via FAO and as guardians against oxidative stress and ferroptotic death position them as major players in PCa growth and progression, warranting additional research into their potential as therapeutic targets.

## MATERIALS AND METHODS

### Cell lines and cell culture

The human prostate cancer cell lines LNCaP, 22Rv1, VCaP and PC3 were obtained from the American Type Culture Collection (ATCC). V16D and 49F^ENZR^ cells^72^ were kindly provided by A. Zoubeidi. LNCaP, 22Rv1, PC3 and 49F^ENZR^ cells were cultured as described previously^73^. V16D cells were maintained in RPMI-1640 containing 10% FBS. VCaP cells were maintained in Dulbecco’s Modified Eagles’s medium (DMEM) containing 10% FBS, 1% sodium pyruvate, 1% MEM non–essential amino acids, and 0.1nM 5α-dihydrotestosterone (DHT). For serum starvation experiments, cells were grown in phenol red-free RPMI-1640 containing 10% dextran-coated charcoal (DCC) stripped serum. Cell Bank Australia performed verification of all cell lines (2016-2020) via short-tandem repeat profiling. All cell lines were subjected to regular mycoplasma testing.

### Cell line transfection

Transfection of cell lines with 10 nM small interfering RNAs (siRNAs) was performed using RNAiMAX Transfection Reagent (Life Technologies) according to the manufacturer’s instructions. SiRNAs used in this study were ACSM1 and ACSM3 Silencer Select (Ambion; siACSM1-1 (s41984), siACSM1-2 (s41985), siACSM3 (s12459) and siACSM3 (s12460)) and control siRNA (Qiazen 1027281).

### Quantitative real-time PCR (qRT-PCR)

Total RNA from cell lines was extracted using TRI Reagent (Sigma), as described previously^74^. Total RNA was treated with Turbo DNA-free kit (Invitrogen), and reverse transcribed using iScript Reverse Transcriptase Supermix kit (Bio-Rad). qRT-PCR was performed in triplicate as described previously^75^. GAPDH levels were used for normalization of qRT-PCR data. Relative gene expression was calculated using the comparative Ct method. Primer sequences are shown in Table 1.

**Table 1.**
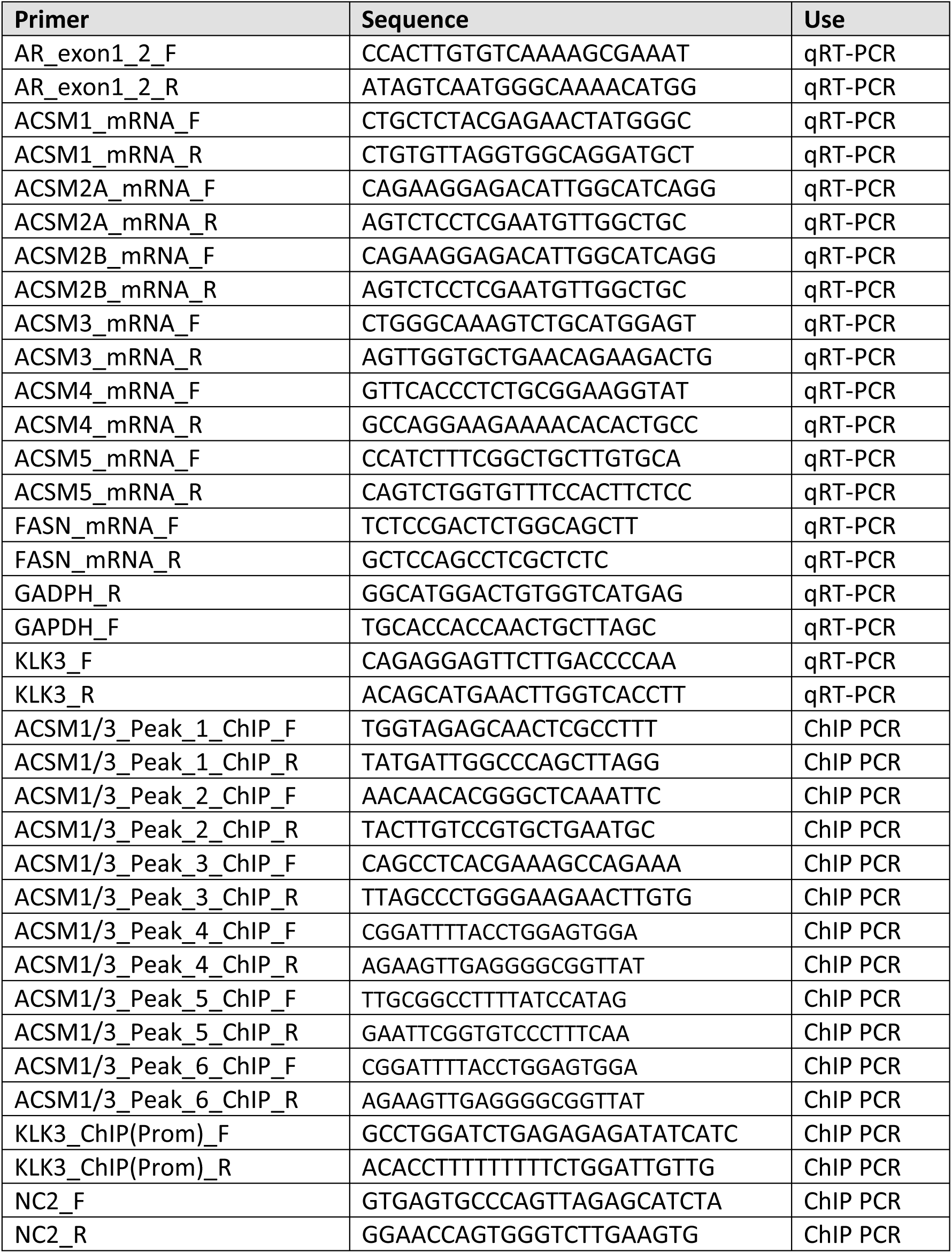
Primers used in this study.

### Chromatin immunoprecipitation (ChIP)

LNCaP cells were seeded at 4 x 10^6^ cells/plate in 15cm plates in RPMI-1640 medium containing 10% DCC-FBS for 3 days, then treated for 4 hours with 10nM DHT or Vehicle (ethanol). AR ChIP using antibody ER179 (Ab108341, Abcam) was performed as described previously^76^.

### Immunoblotting

Protein extraction from cells using RIPA buffer and western blotting was done as described previously^75^. The following primary antibodies were used: AR (N20, SC-816, Santa Cruz Biotechnology); β-actin (AC-15, Ab6376, Abcam); ACSM1 (SAB1301678, Sigma-Aldrich); ACSM3 (sc-377173 (G8), Santa Cruz Biotechnology); DLAT (12362S, Cell Signalling Tech); DLST (11954S, Cell Signalling Tech); Lipoic acid (ab58724, Abcam); α-tubulin (05-829, Millipore); GAPDH (MAB374, Millipore); MDA (ab6463, Abcam); and GPX4 (ab41787, Abcam).

### Analysis of published RNA-seq data

Clinical transcriptomic data was downloaded from GEO (GSE7868^14^; GSE22606^15^; and GSE51005^21^) and cBioportal (MSKCC^16^; TCGA^77^; SU2C^78^). CPGEA data^79^ was obtained from http://bigd.big.ac.cn/gsa-human/.

### Immunohistochemistry (IHC)

IHC was performed essentially as described previously^80^. Tissue sections were incubated with primary antibodies ACSM1 (1:100, SAB1301678, Sigma-Aldrich) and ACSM3 (1:50, sc-377173 (G8), Santa Cruz Biotechnology) overnight at 4°C. Biotinylated secondary antibodies (E0432 and E0433, Dako) were added to slides at a 1:400 dilution in blocking solution for 1h at RT followed by HRP-streptavidin at 1:500 for 1 hr at RT. Slides were scanned using a Nanozoomer digital slide scanner (Hamamatsu). Images were viewed with NDP view software and quantified using ALLRED score^81^.

### Proliferation assays

Proliferation of cells transfected with siACSM1 and siACSM3 was measured using Trypan blue exclusion assays. Cells were seeded at 6 × 10^4^ (LNCaP, 49F^ENZR^, 22RV1), 1 × 10^5^ (VCaP) and 1 × 10^4^ (PC3) in 12-well plates. Live cells were quantified using Trypan blue.

### Apoptosis assays

Apoptosis was measured by collecting cells in FACS binding buffer (47 ml of HANKS buffered saline, 500 µL of HEPES solution and 2.5 mL of 100 mM CaCl2), staining with Annexin V PE (BD Biosciences) and 1 mM 7-Aminoactiomycin D (Thermo Fisher Scientific) and analysis by Flow Cytometry using a LSRFortessa X20 (BD Biosciences).

### Colony and spheroid formation assays

For colony formation assays, LNCaP cells were washed with PBS, trypsinized, collected, counted and single-cell suspension was prepared. 500 cells were plated in 6-well plates and incubated for 2 weeks at 37°C, with media replenishment every 3-7 days. Cells were then washed with PBS, fixed with 4% paraformaldehyde and stained with 1% crystal violet for 30 minutes. Colonies were counted manually.

For spheroid assays, LNCaP cells were collected and prepared at a concentration of 7.5 × 10^4^ cells/ml. Cell suspensions (1500 cells in 20 µl) were pipetted onto 96 well Spheroid ULA/CS plates with 100 µL of media and incubated at 37°C for 5 days. Photos of the formed spheres were captured and sphere volume was determined using ReViSP software^82^.

### Reactive oxygen species (ROS) assays

Cellular ROS levels were measured using CellROX^TM^ Orange Flow Cytometry Assay Kits (Life Technologies) as described previously^83^. MitoSOX Red, a mitochondrial superoxide indicator, was measured by flow cytometry as described previously^31^.

### Lipid analysis by imaging

LNCaP cells were seeded in an 8-well chamber slide (5 × 10^3^ cells/well) and transfected with siACSM1, siACSM3 or siNC for 96 h. Cells were stained with 5 μM BODIPY-581/591 C11 (Thermo Fisher Scientific) as described previously^31^. Quantification of BODIPY-C11 stain was performed using ImageJ analysis software as described previously^31^.

### Lipid analysis by flow cytometry

To measure total lipid droplets, LNCaP cells (2 x 10^5^ cells/well) were transfected with siACSM1, siACSM3 or siNC for 48 h in 6 well plates. Alternatively, over-expression lines (ACSM1-OE, ACSM3-OE or control) cells (3 x 10^5^ cells/well) were seeded in 6 well plate for 48 hours. Cells were harvested by trypsinizing, collected in FACS tubes, centrifuged at 450 g for 5 mins and then washed with FACS binding buffer. Cells were resuspended in 0.2 ml of BODIPY 493/503 (2 µM) in PBS and incubated at 37°C for 15 min in the dark. After carefully aspirating the stain, cells were washed again with PBS, pelleted by centrifugation at 450 g for 5 mins and resuspended in 0.2 ml of FACS binding buffer. Flow cytometric analysis was performed on an LSRFortessa X20 (minimum 10,000 events). Three independent biological replicates were performed for each condition. For total lipid analysis experiments, oleic acid (200 µM in 2% BSA) was added to cells 24 h prior to staining as a positive control.

To measure lipid peroxidation, LNCaP cells (2 x 10^5^ cells/well) were transfected with siACSM1, siACSM3 or siNC for 48 h in 6 well plates. Cells were harvested by trypsinizing, collected in FACS tubes, centrifuged at 450 g for 5 mins and then washed with Hank’s balanced salt solution (HBSS). Cells were resuspended in 0.2 ml of C11 BODIPY 581/591 (5 µM) dissolved in HBSS and incubated at 37°C for 10 min. After carefully aspirating the stain, cells were pelleted by centrifugation at 450 g for 5 mins and resuspended in 0.2 ml fresh HBSS. Flow cytometric analysis was performed on an LSRFortessa X20. Oxidation of C11 BODIPY 581/591 was calculated as the ratio of the green fluorescence (indicating oxidized probe) to total (green + red) fluorescence (indicating reduced plus oxidized probe). Three independent biological replicates were performed for each condition. For lipid peroxidation experiments, Erastin (10 uM) was used as a positive control.

### ATP assays

LNCaP cells were transfected with siACSM1 and siACSM3 for 72 hours in 96 well plates. Total cellular ATP was measured using a Luminescent ATP Detection Assay Kit (Abcam), according to the manufacturer’s instructions. ATP measurements were normalized to the total cellular DNA content in each well, which was determined using a CyQUANT Cell Proliferation Assay kit (ThermoFisher Scientific).

### Seahorse extracellular flux analysis

Cells were plated in XF96 well cell culture microplates (Agilent) at equal densities in substrate-limited medium (DMEM with 0.5 mM glucose, 1.0 mM glutamine, 0.5 mM carnitine and 1% FBS) and incubated overnight. One hour before the beginning OCR measurements, the cells were changed into FAO Assay Medium (111 mM NaCl, 4.7 mM KCl, 2.0 mM MgSO_4_, 1.2 mM Na_2_HPO_4_, 2.5 mM glucose, 0.5 mM carnitine and 5 mM HEPES). After baseline OCR was stabilized in FAO Assay Medium, 200 µM of linoleic-acid (LA) or palmitic acid (PA) were added before initializing measurements. Extracellular flux analysis was performed using the Seahorse XF Cell Mitochondrial Stress Test kit (Seahorse Bioscience) according to the manufacturer’s protocol. Extracellular flux experiments were performed on a Seahorse XF96 Analyzer and results were analysed using Seahorse Wave software for XF analyzers. The OCR values were normalized to cell numbers in each well.

### Glutathione assays

Cells were seeded in in poly L-lysine coated 6-well plates (1.5 × 10^5^ cells/well), transfected with 10nM of siACSM1, siACSM3 or siNC and allowed to grow for 7 days. The cells were collected by scraping and glutathione (GSH+GSSG) was measured using a Cayman Chemical Glutathione Assay Kit (Cayman Chemical), according to the manufacturer’s protocol. GSH and GSSG concentrations were calculated using a standard curve and normalized to the total protein level in each sample. Three independent biological replicates were performed for each condition.

### Lipidomics

LNCaP cells (2 x 10^5^ cells/well in 6 well plates) were transfected with siACSM1/3 for 4 or 6 days. Alternatively, over-expression lines (ACSM1-OE, ACSM3-OE or control) were seeded in 6 well plates (2 x 10^5^ cells/well) and grown for 4 days. At the end of the treatment/growth period, culture plates were placed on ice. Media was removed and cells were washed 3 times with cold PBS, after which they were scraped and collected in 1.5 ml tubes by centrifugation at 20,000g for 5 min. The supernatant was removed and cell pellets were frozen and stored at -80°C. To extract lipids, 700 μl of sample (4 μl of plasma diluted in water, or 700 μl of homogenized cells) was mixed with 800 μl 1 N HCl:CH3OH 1:8 (v/v), 900 μl CHCl3 and 200 μg/ml of the antioxidant 2,6-di-tert-butyl-4-methylphenol (BHT; Sigma Aldrich). 3 μl of SPLASH LIPIDOMIX Mass Spec Standard (#330707, Avanti Polar Lipids) was spiked into the extract mix. The organic fraction was evaporated using a Savant Speedvac spd111v (Thermo Fisher Scientific) at room temperature and the remaining lipid pellet was stored at -20°C under argon.

Lipid pellets were reconstituted in 100% ethanol. Lipid species were analyzed by liquid chromatography electrospray ionization tandem mass spectrometry (LC-ESI/MS/MS) on a Nexera X2 UHPLC system (Shimadzu) coupled with hybrid triple quadrupole/linear ion trap mass spectrometer (6500+ QTRAP system; AB SCIEX). Chromatographic separation was performed on a XBridge amide column (150 mm × 4.6 mm, 3.5 μm; Waters) maintained at 35°C using mobile phase A [1 mM ammonium acetate in water-acetonitrile 5:95 (v/v)] and mobile phase B [1 mM ammonium acetate in water-acetonitrile 50:50 (v/v)] in the following gradient: (0–6 min: 0% B → 6% B; 6–10 min: 6% B → 25% B; 10–11 min: 25% B → 98% B; 11–13 min: 98% B → 100% B; 13–19 min: 100% B; 19–24 min: 0% B) at a flow rate of 0.7 mL/min which was increased to 1.5 mL/min from 13 min onwards. SM, CE, CER, DCER, HCER, LCER were measured in positive ion mode with a precursor scan of 184.1, 369.4, 264.4, 266.4, 264.4 and 264.4 respectively. TAG, DAG and MAG were measured in positive ion mode with a neutral loss scan for one of the fatty acyl moieties. PC, LPC, PE, LPE, PG, LPG, PI, LPI, PS and LPS were measured in negative ion mode by fatty acyl fragment ions. Lipid quantification was performed by scheduled multiple reactions monitoring (MRM), the transitions being based on the neutral losses or the typical product ions as described above. The instrument parameters were as follows: Curtain Gas = 35 psi; Collision Gas = 8 a.u. (medium); IonSpray Voltage = 5500 V and −4,500 V; Temperature = 550°C; Ion Source Gas 1 = 50 psi; Ion Source Gas 2 = 60 psi; Declustering Potential = 60 V and −80 V; Entrance Potential = 10 V and −10 V; Collision Cell Exit Potential = 15 V and −15 V. The following fatty acyl moieties were taken into account for the lipidomic analysis: 14:0, 14:1, 16:0, 16:1, 16:2, 18:0, 18:1, 18:2, 18:3, 20:0, 20:1, 20:2, 20:3, 20:4, 20:5, 22:0, 22:1, 22:2, 22:4, 22:5 and 22:6 except for TGs which considered: 16:0, 16:1, 18:0, 18:1, 18:2, 18:3, 20:3, 20:4, 20:5, 22:2, 22:3, 22:4, 22:5, 22:6.

Peak integration was performed with the MultiQuant software version 3.0.3. Lipid species signals were corrected for isotopic contributions (calculated with Python Molmass 2019.1.1) and were normalized to internal standard signals. Unpaired T-test p-values and FDR corrected p-values (using the Benjamini/Hochberg procedure) were calculated in Python StatsModels version 0.10.1.

### Metabolomics

LNCaP cells (5.0 x 10^6^) were transfected with siRNA for 48 hours in RPMI (phenol red free) with 10% FBS supplemented with 2.0 mM glucose in 6-well plates. Cells were placed on ice and washed twice with 5 ml cold NaCl saline, after which 300µl of ice cold methanol:chloroform (MeOH:CHCl3) extraction solvent containing the internal standards (0.5µl/samples) was added onto each well. Cells were scraped and collected in 15ml tubes, after which an additional 300 µL MeOH:CHCl3 was added to wells in order to collect remaining cells. 1.5 ml of chloroform was added to each tube followed by vortexing and incubation on ice for 5 mins. Tubes were centrifuged at 4°C for 5 minutes at 2,700g and the top aqueous layer was then transferred into a fresh 1.5ml Eppendorf tube and allowed to dry in a Speedvac. Dried samples were derivatised with 20 µl methoxyamine (30 mg/ml in pyridine, Sigma Aldrich) and 20 µl N,O-Bis(trimethylsilyl)trifluoroacetamide (BSTFA) + 1% Trimethylchlorosilane (TMCS). The derivatised samples were analysed using GC QQQ targeted metabolomics as described^84^.

### Lentiviral transduction of cells

SMARTvector Inducible Lentiviral shRNA vectors for ACSM1, ACSM3 and Non-targeting shRNA Control (glycerol stock *E. coli*) were obtained from Dharmacon (catalogue numbers V3SH11252-225482707 and V3SH11252-224860261). For over-expression of ACSM1 and ACSM3 in PCa cells, linear dsDNA fragments containing codon-optimised ACSM1 and ACSM3 cDNA sequences were cloned into Gateway entry vector (pDONR221) and then transferred into pJS64 destination vector. The sequence of the inserts was verified by Sanger sequencing at AGRF. Viral supernatants were prepared by transfection of HEK293T cells with the vector plasmids and packaging plasmids (psPAX2 Addgene 12259 and pMD2.G Addgene 12260). The transfection was performed with polyethylenimine (PEI MAX Transfection Grade Linear Polyethylenimine Hydrochloride, 24765-1, Polysciences) as described previously^74^. After 48 hours, supernatants were filtered through 0.45 µm and 0.22 µm syringe filters and then concentrated ∼100-fold using Vivaspin 100 kDa cut-off columns by centrifugation at 3000 g at 10°C for 6 hours. Concentrated virus was stored at -80°C. For transduction, luciferase-tagged LNCaP cells (for shRNA viruses) and untagged LNCaP cells (for over-expression vectors) were seeded at 1.25 × 10^5^ cells in 6 well plates and the following day were transduced with concentrated lentivirus for a period of 3 days. Puromycin (1.0 μg per mL) was added to select for transduced cells for an additional 3 days. Cells transduced with shRNA constructs were grown in 2ug µg per ml of doxycycline to induce expression of shRNAs.

### Animal experiments

All animal procedures were approved by the University of Adelaide Animal Ethics Committee (approval number M-2019–037) and carried out in accordance with the guidelines of the National Health and Medical Research Council of Australia. LNCaP-NC and LNCaP-shACSM3 cell suspensions (1 × 10^6^ cells in 10 µl PBS) were injected intraprostatically in 8-week old NOD scid gamma male mice. After 10 days, mice were fed drinking water containing doxycycline (2mg/mL) and sucralose (1 mg/mL; added as a sweetener). Doxycycline-treated water was freshly prepared and changed weekly in light-protected bottles until the study was terminated. Whole-body imaging to monitor luciferase-expressing LNCaP cells was performed 3 days after injection of tumour cells and once weekly thereafter using an IVIS Spectrum *in vivo* Imaging System (PerkinElmer). D-luciferin (potassium salt, PerkinElmer) was dissolved in sterile deionized water (0.03 g/ml) and injected subcutaneously (3 mg/20 g of mouse body weight) before imaging. Bioluminescence was reported as the sum of detected photons per second from a constant region of interest.

### Statistical analyses

All experiments were performed at least three times. Statistical analysis was performed using GraphPad Prism 7.02 software. Statistical methods are included in figure legends.

## ACKNOWLEDGEMENTS

This work was supported by: the Movember Foundation and the Prostate Cancer Foundation of Australia through a Movember Revolutionary Team Award (MRTA-3 to LMB, AJH, WDT and LAS); the National Health and Medical Research Council of Australia (1121057 to WDT and LAS); Cancer Council SA (Project Grant 1185012 to LAS); and The Hospital Research Foundation (Project Grant C-PJ-126-Exper-2019 to LAS). LAS and LMB are supported by Principal Cancer Research Fellowships (PRF2919 and PRF1117, respectively) awarded by Cancer Council’s Beat Cancer project on behalf of its donors, the state Government through the Department of Health and the Australian Government through the Medical Research Future Fund.

The authors thank: Ge Liu and Robert Gibson for assistance with lipid analysis; Randall Grose for expert technical assistance with flow cytometry; and Joanna Gillis, Natalie Ryan, Geraldine Laven-Law, Zoya Kikhtyak and Rayzel Fernandes for expert technical assistance. The results published here are in part based on data generated by The Cancer Genome Atlas, established by the National Cancer Institute and the National Human Genome Research Institute, and we are grateful to the specimen donors and relevant research groups.

## AUTHOR CONTRIBUTIONS

RS, LMB and LAS conceived the project. RS, JVS, LMB and LAS designed experiments. RS, ZDN, ARH, SLT, JD, CYM, LEQ and MP performed experiments and acquired data. SLT, ARH, RI, CYM, MH, MG, LEQ and MP provided technical assistance. RS, ZDN, ARH, RI, SLT, JD, CYM, MH, MG, MP, MJW, LEQ, AJH, WDT, JVS, LMB and LAS interpreted and analysed data. RS, LMB and LAS wrote the manuscript. All of the authors have read, edited, and approved the paper.

## DECLARATION OF INTERESTS

The authors have no conflicts of interest to disclose.

---

**Supplementary Figure S1.**
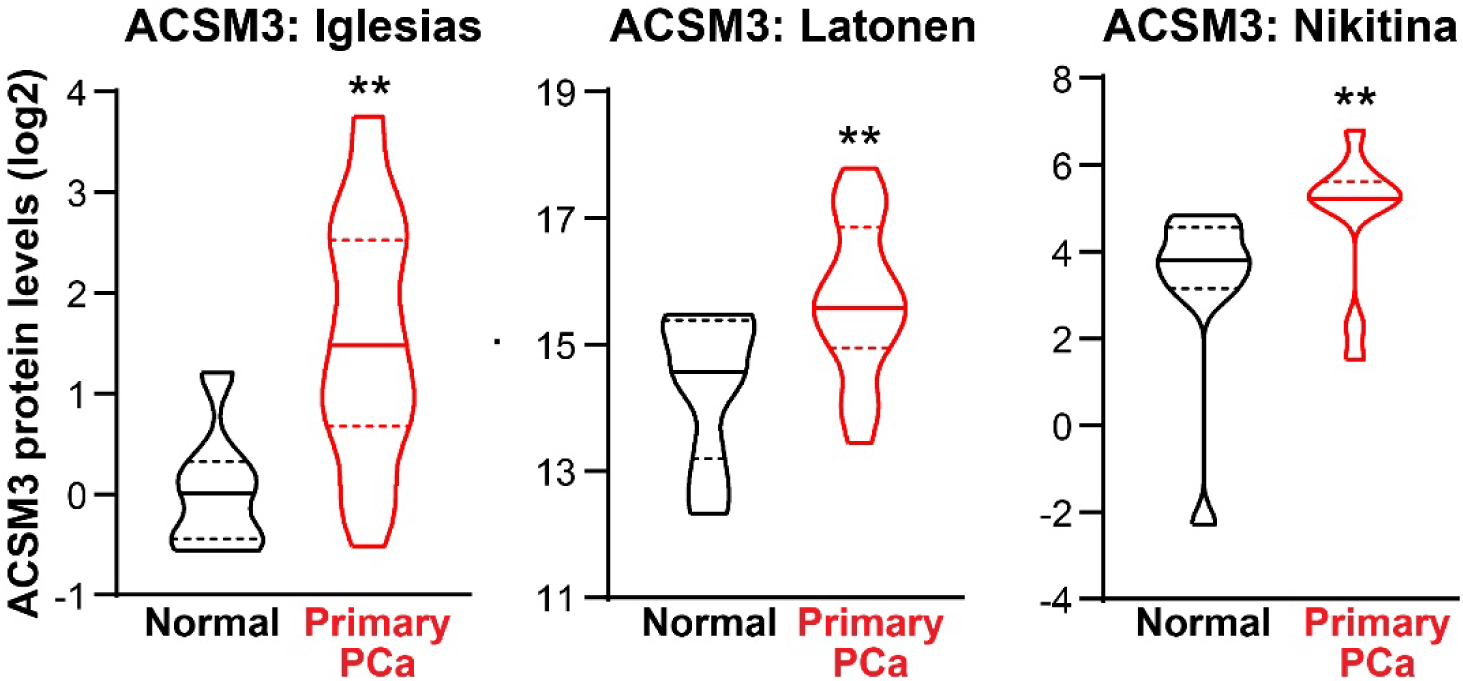
ACSM3 protein levels are elevated in primary prostate cancer (PCa) compared to normal prostate tissues. Violin plots show minimum and maximum (bottom and top lines, respectively) and mean (line within the boxes) values. Unpaired t tests were used to compare expression in normal and cancer tissues (**, p < 0.01).

**Supplementary Figure S2.**
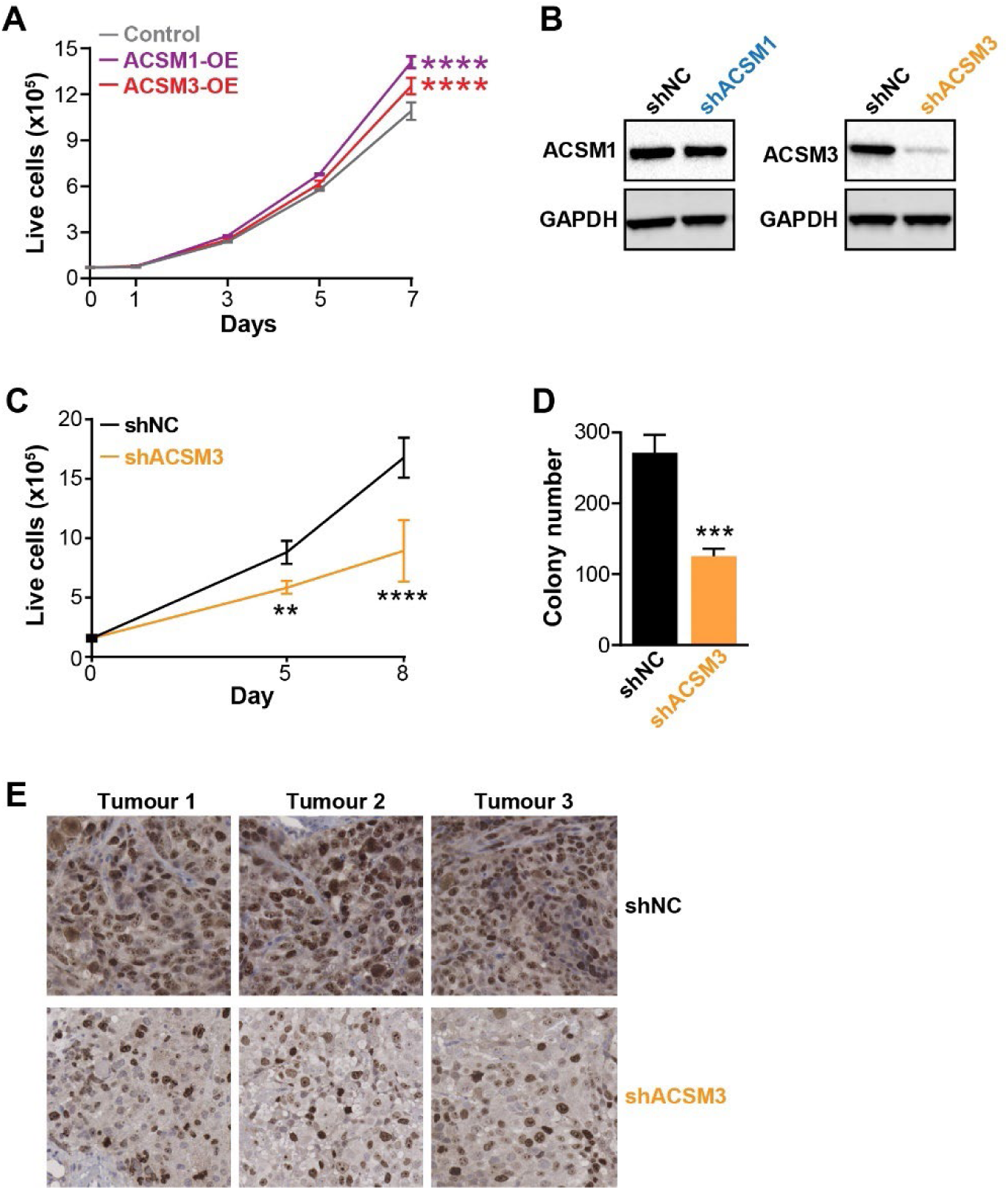
**(A)** Over-expression of ACSM1 and ACSM3 promotes growth of LNCaP cells, as evaluated by trypan blue growth assays. Growth was compared to control cells (at day 7) using ANOVA and Dunnett’s multiple comparison tests. **(B)** ACSM1 and ACSM3 protein levels in LNCaP derivatives stably transduced with shACSM1 and shACSM3. Cells were cultured in doxycycline (2 µg/mL) containing normal growth media for 5 days after which ACSM1 and ACSM3 proteins were evaluated by immunoblotting. GAPDH was used as a loading control. **(C)** Trypan blue assay showing growth of shNC and shACSM3 cells grown in the presence of doxycycline (2 µg/mL). Unpaired t tests were used to compare growth at the indicated time-points. **(D)** Clonogenic cell survival of LNCaP-shACSM3 cells was assessed using a colony formation assay. Cells were cultured for 2 weeks in normal growth media containing doxycycline (2 µg/mL). Colonies were visualised and counted as for Figure 3D. Data shown is representative of n = 3 independent experiments. An unpaired t test was used to compare shNC and shACSM3. For all statistical tests: **, p < 0.01; ***, p < 0.001; ****, p < 0.0001. **(E)** Representative Ki67 immunohistochemistry images of Ki67 positivity in LNCaP orthotopic tumours collected at day 45. Ten such representative images per tumour, collected at ×20 magnification, were used to quantify Ki67 positivity in each condition (Figure 3K).

**Supplementary Figure S3.**
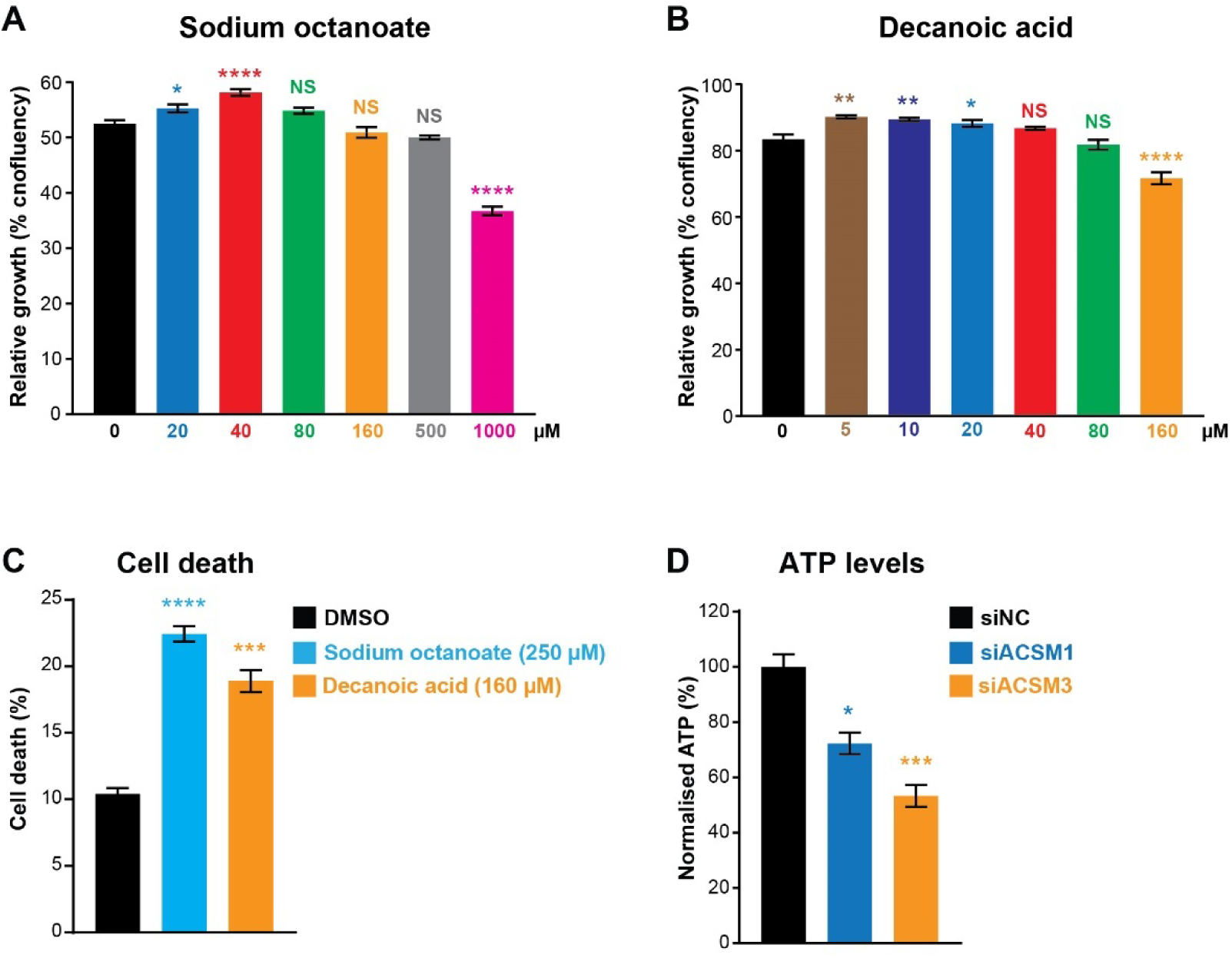
**(A-B)** Effect of sodium octanoate (A) and decanoic acid (B) on growth of LNCaP cells. Graphs represent relative cell confluency (114 hours after seeding), measured using an Incucyte. One way ANOVA and Tukey’s multiple comparisons test were used to compare groups. **(C)** Effect of growth-inhibitory doses of sodium octanoate and decanoic acid on LNCaP cell death, as determined using flow cytometry-based 7-AAD assays. Data represents the mean ± SEM of triplicate samples and are representative of 4 independent experiments. One way ANOVA and Dunnett’s multiple comparisons test were used to compare groups. **(D)** Effect of ACSM1 and ACSM3 knockdown on ATP levels in LNCaP cells. Data was normalised to control (siNC), which was set to 100%. Unpaired t tests were used to compare siACSM1/siACSM3 relative to siNC. For all statistical tests: *, p < 0.05; **, p < 0.01; ***, p < 0.001; ****, p < 0.0001.

**Supplementary Figure S4.**
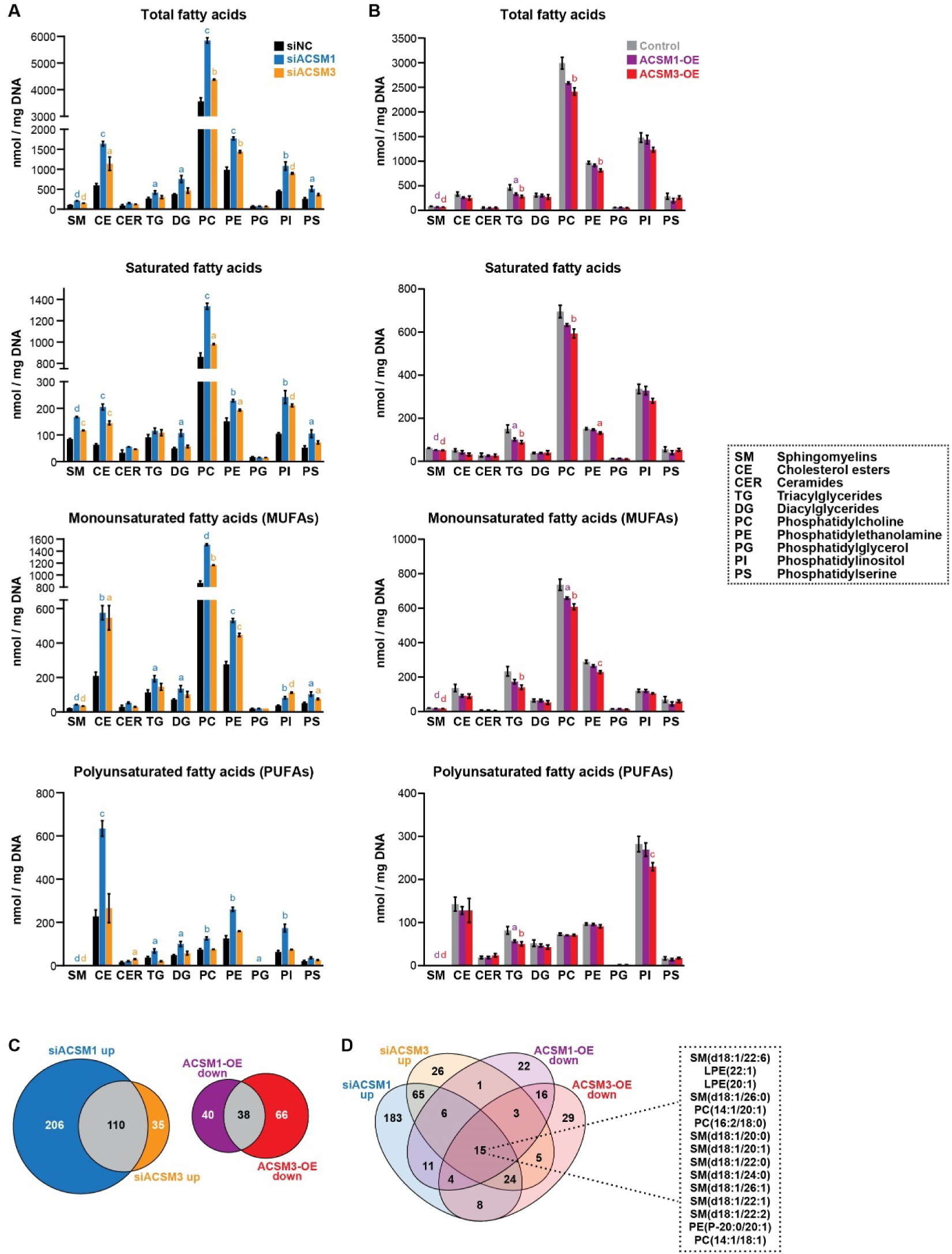
ACSM1 and ACSM3 are major regulators of the prostate cancer lipidome. **(A)** Abundance of lipid species in response to knockdown of ACSM1 and ACSM3. Top graph shows total lipids in each class; bottom 3 graphs show lipids according to saturation status. Data are mean ± SEM. Unpaired t tests were used to compare siACSM1/siACSM3 relative to siNC. **(B)** Abundance of lipid species in response to over-expression of ACSM1 and ACSM3. Data is presented as in (A). For all statistical tests: a, p < 0.05; b, p < 0.01; c, p < 0.001; d, p < 0.0001. **(C)** Venn diagrams showing overlap of lipids upregulated in response to siACSM1 or siACSM3 (left) or lipids downregulated in response to over-expression of ACSM1 or ACSM3 (right). **(D)** Venn diagram showing overlap of lipids upregulated in response to siACSM1/siACSM3 or lipids downregulated in response to over-expression of ACSM1/ACSM3. The 15 lipid species altered in each experimental condition are shown.

**Supplementary Figure S5.**
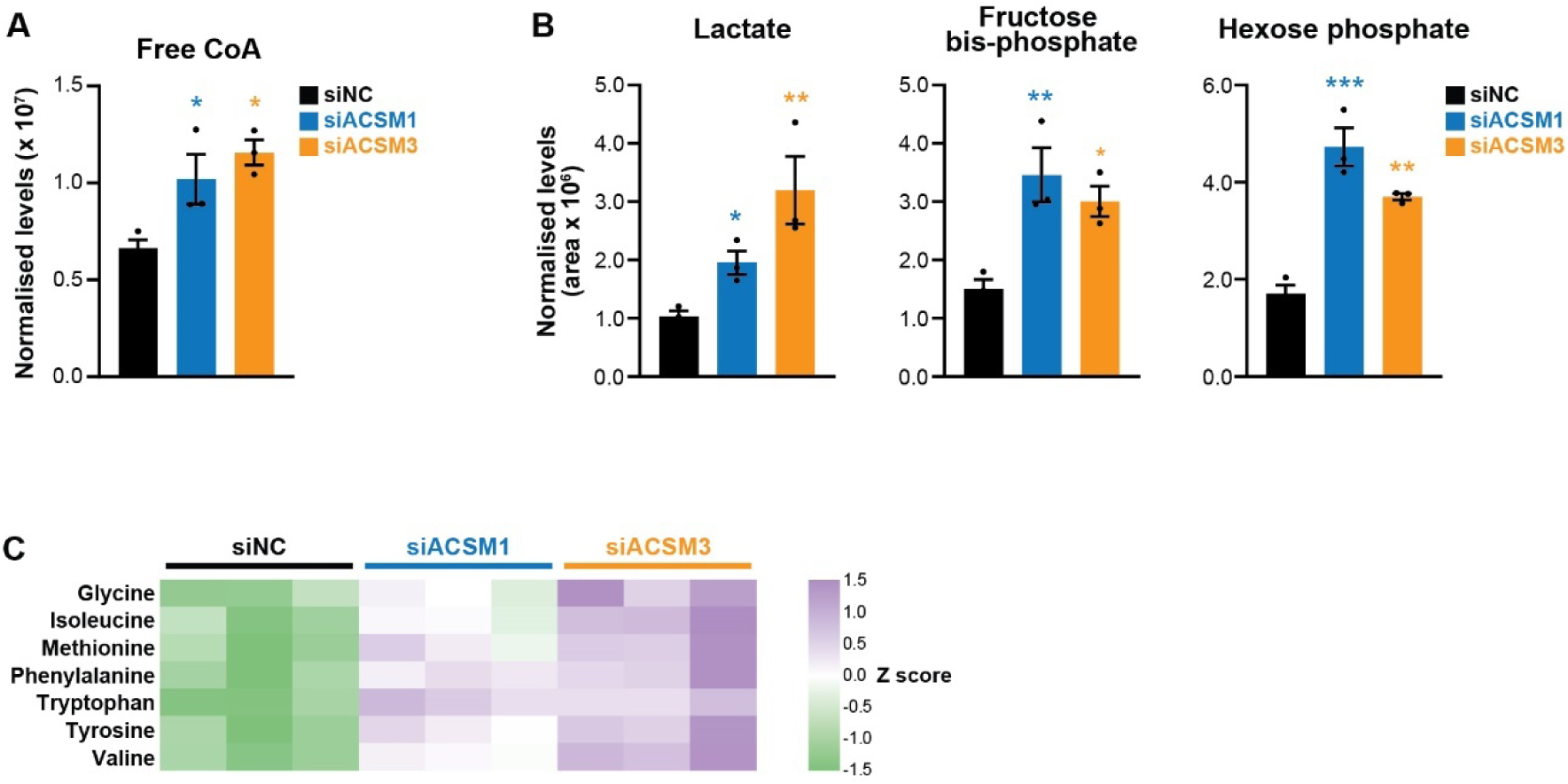
**(A-B)** Loss of ACSM1 and ACSM3 in LNCaP cells causes accumulation of free CoA (A), lactate (B) and glycolytic intermediates (B), measured using GC QQQ targeted metabolomics. Data in bar graphs are mean ± SEM. Unpaired t tests were used to compare siACSM1/siACSM3 relative to siNC (*, p < 0.05; **, p < 0.01; ***, p < 0.001; ****, p < 0.0001). **(C)** Loss of ACSM1 and ACSM3 results in accumulation of amino acids, measured using GC QQQ targeted metabolomics. Z scores were derived from metabolomic data; 3 replicates of each treatment (siNC, siACSM1 and siACSM3) are shown.

**Supplementary Figure S6.**
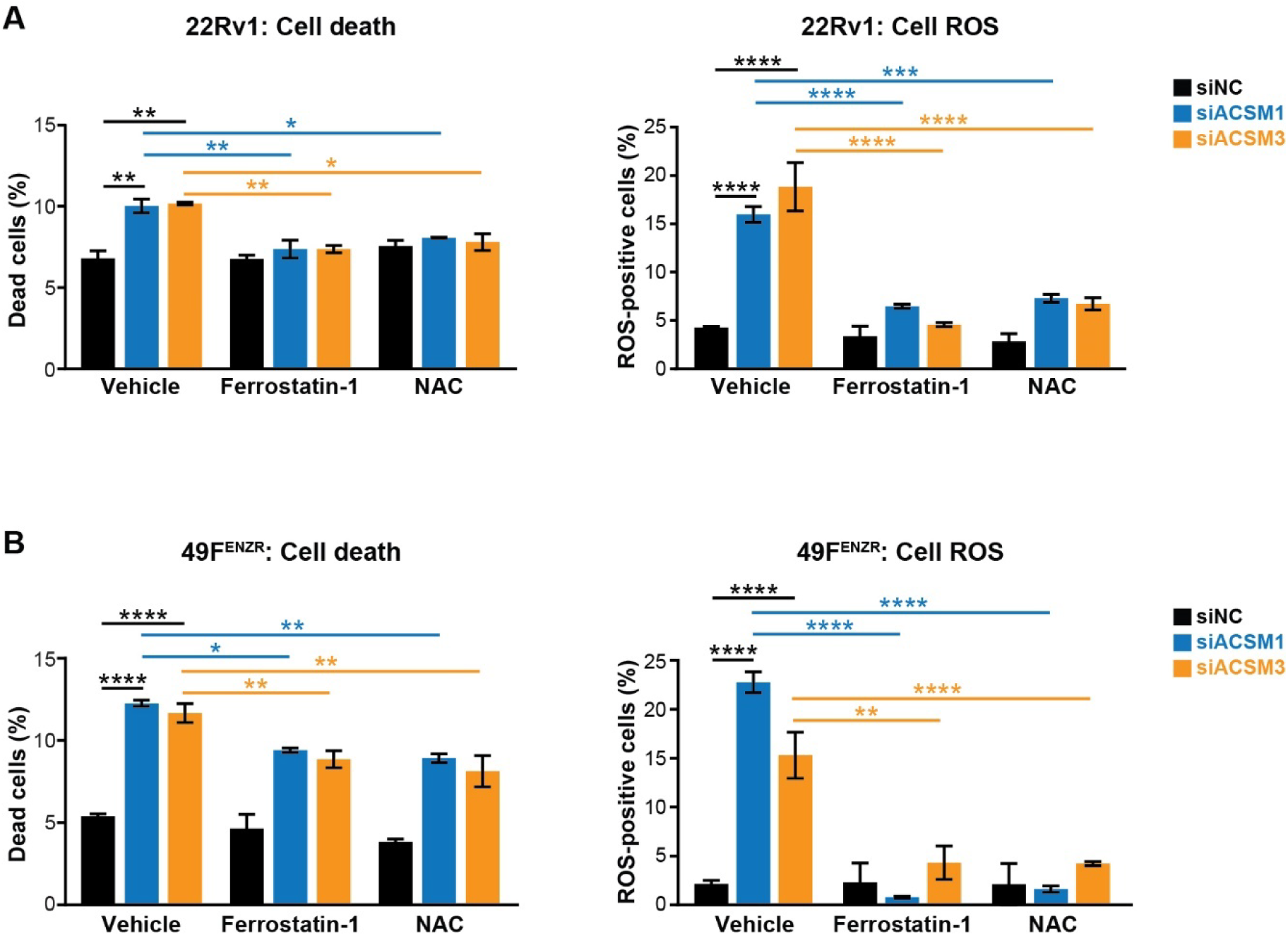
**(A-B)** Ferrostatin-1 and NAC rescue cell death and ROS induction mediated by loss of ACSM1 and ACSM3 in 22Rv1 (A) and 49F^ENZR^ (B) cells. Data are mean ± SEM. One way ANOVA and Tukey’s multiple comparisons test were used to compare groups.

## Notes

### Competing Interest Statement

The authors have declared no competing interest.

